# Using GPUs to accelerate computational diffusion MRI: From microstructure estimation to tractography and connectomes

**DOI:** 10.1101/371278

**Authors:** Moises Hernandez-Fernandez, Istvan Reguly, Saad Jbabdi, Mike Giles, Stephen Smith, Stamatios N. Sotiropoulos

## Abstract

The great potential of computational diffusion MRI (dMRI) relies on indirect inference of tissue microstructure and brain connections, since modelling and tractography frameworks map diffusion measurements to neuroanatomical features. This mapping however can be computationally highly expensive, particularly given the trend of increasing dataset sizes and the complexity in biophysical modelling. Limitations on computing resources can restrict data exploration and methodology development. A step forward is to take advantage of the computational power offered by recent parallel computing architectures, especially Graphics Processing Units (GPUs). GPUs are massive parallel processors that offer trillions of floating point operations per second, and have made possible the solution of computationally-intensive scientific problems that were intractable before. However, they are not inherently suited for all problems. Here, we present two different frameworks for accelerating dMRI computations using GPUs that cover the most typical dMRI applications: a framework for performing biophysical modelling and microstructure estimation, and a second framework for performing tractography and long-range connectivity estimation. The former provides a front-end and automatically generates a GPU executable file from a user-specified biophysical model, allowing accelerated non-linear model fitting in both deterministic and stochastic ways (Bayesian inference). The latter performs probabilistic tractography, it can generate whole-brain connectomes and supports new functionality for imposing anatomical constraints, such as inherent consideration of surface meshes (GIFTI files) along with volumetric images. We validate the frameworks against well-established CPU-based implementations and we show that despite the very different challenges for parallelising these problems, GPU-based designs can offer accelerations of more than two orders of magnitude in both cases.

## INTRODUCTION

General-purpose computing on graphics processing units (GPGPU) has lead to a significant step forward in scientific computations. GPUs are massive parallel processors with thousands of cores. Mainly driven by the computer game industry, and more recently by deep learning applications (Schmidhuber 2015), GPUs have evolved rapidly in the last decade, offering now over 15 TeraFLOPS (15×10^13^ floating operations per second) in single precision of performance (NVIDIA 2017). Even if their full potential is not used, their suitability for scientific computing has become more and more evident in projects that involve large amounts of data. For instance, the 1000 Genomes Project (Auton et al. 2015; Sudmant et al. 2015) and the Human Connectome Project (Van Essen & Ugurbil 2012; Van Essen et al. 2012; Sotiropoulos et al. 2013) have generated Petabytes of data. The computations performed for the analysis of all this data can take months on typical computer clusters, but GPU accelerated solutions can accelerate massively these computations (Hernández et al. 2013; Klus et al. 2012).

In the field of medical imaging, GPUs have been used in the last decade in several computational domains (Eklund et al. 2013), including image reconstruction (Stone et al. 2008; Uecker et al. 2015) image segmentation (Alsmirat et al. 2017; Smistad et al. 2015), image registration (Shamonin 2013), and in the analysis of functional MRI (Eklund et al. 2014) and diffusion MRI data(Xu et al. 2012; Hernández et al. 2013; Chang et al. 2014; Hernandez-Fernandez et al. 2016; Harms et al. 2017).

However, using GPUs is not always straightforward. The GPU architecture is completely different to the traditional single or multi-core CPU architectures, it is not inherently suited for all types of problems, and bespoke computational frameworks may need to be developed to take advantage of their full potential. Some of the challenges that need to be considered for achieving an efficient design include: balanced parallelisation of an algorithm, good organisation of threads and grouping, appropriate usage of memory resources, appropriate memory access patterns, and correct communication and synchronisation between threads. Furthermore, programming GPUs requires specific programming models that offer control over the device resources, but may increase the difficulty for designing parallel solutions. Low-level programming models, such as the Compute Unified Device Architecture (CUDA) (Nickolls et al. 2008), offer a high degree of control over the resources, and the possibility of achieving very efficient solutions.

Despite the challenges, in this paper we illustrate the potential of GPUs for two neuroimaging applications spanning different parallelisation strategies. Specifically, we design and implement parallel computational frameworks for analysing diffusion magnetic resonance imaging (dMRI) data (Sotiropoulos et al. 2013; Johansen-Berg & Behrens 2014). The great potential of dMRI is that it uniquely allows studying the human brain non-invasively and in vivo. However, it relies on indirect inference from the data, and typically, modelling frameworks are necessary to map dMRI measurements to neuroanatomical features, which can be computationally expensive. The computational cost is becoming even higher given the trend of increasing data sizes. New MRI hardware and sequences for pushing spatial and angular resolution (Vu et al. 2015; Setsompop et al. 2017) can considerably increase the size of a single subject dataset. At the same time, big imaging data repositories with a large number of subjects are being created, such as the Human Connectome Project (HCP) (Van Essen & Ugurbil 2012; Van Essen et al. 2012; Sotiropoulos et al. 2013), where 1,200 datasets are included, its Lifespan and Disease extensions (https://www.humanconnectome.org) and UK Biobank (Miller et al. 2016; Alfaro-Almagro et al. n.d.) where a total of 100,000 subjects are being scanned. Limitations in computing can restrict data exploration and even methodology development.

A common application of dMRI analysis is tissue microstructure estimation. Even if the diffusion tensor imaging (DTI) (P. Basser et al. 1994; P. J. Basser et al. 1994) model is by far the most popular framework for extracting microstructure indices and can be fitted linearly to data, it has major limitations, such as the inability to capture partial volume, leading to non-specific markers of tissue structural changes (Basser et al. 2000; Pierpaoli et al. 2001; Poupon et al. 2000; Wiegell et al. 2000; Alexander et al. 2001; Seunarine & Alexander 2014). To overcome these limitations multi-compartment biophysical models are being developed, where the diffusion signal attenuation is represented as a mixture of signals obtained from different tissue components (Assaf et al. 2008; Alexander et al. 2010; Zhang et al. 2012; Sotiropoulos et al. 2012; Szafer et al. 1995; Niendorf et al. 1996; Stanisz et al. 1997; Assaf & Cohen 1998; Mulkern et al. 1999). Multi-compartment dMRI models are commonly non-linear functions of the signal, and non-linear optimisation algorithms are typically used for fitting the model to the diffusion-weighted measurements (Motulsky & Ransnas 1987; Kelley 1999). These algorithms use iterative optimisation procedures for finding a global solution, leading to potentially large computational times. Furthermore, if a stochastic (instead of deterministic) optimisation method is required, such as Bayesian approaches (Tarantola 2005), computation times are even longer. For instance, using a single CPU core, fitting the NODDI-Bingham model (Tariq et al. 2016) to a single subject UK Biobank dataset using the current available Matlab toolbox (Microstructure Imaging Group - University College London n.d.) requires more than a week for deterministic fitting, and fitting the ball and sticks model (Behrens et al. 2003; Behrens et al. 2007) to a single subject HCP dataset using FMRIB’s Software Library (FSL) (Jenkinson et al. 2012) requires more than 4 days for stochastic fitting using MCMC (see Table 1).

**Table 1.** Execution times and memory requirements of some dMRI applications processing datasets from the UK Biobank project and the Human Connectome Project. Processing times are reported using a single CPU core of a modern processor (Intel Xeon E5-2660 v3 processor).

| **Application (Single subject)** | **Single CPU core time** | **Memory Required** |
| --- | --- | --- |
| Ball & sticks model (MCMC) - UK Biobank | 4 days | 2.5 GB |
| Ball & sticks model (MCMC) - HCP | 11 days | 8 GB |
| NODDI-Bingham. Multi-compartment model - HCP | 7.5 days | 2.5 GB |
| Brain Connectome (dense) - HCP | 15 days | 35 GB |

Optimisation methods used for fitting voxel-wise biophysical models to dMRI data are inherently parallelisable and in general well suited for GPU design, since the computational modelling is applied independently to different voxels. The large number of independent elements in the data and the fact that identical procedures need to be performed over each of these elements make GPUs a perfect candidate for processing these datasets, as they have been designed for exploiting the data level parallelism by executing the same instructions over different data simultaneously (SIMD (Flynn 1972)). However, due to the heavy tasks involved in the optimisation procedures, the design of an optimal parallel solution is non-trivial.

A number of GPU frameworks for accelerating these routines have been developed in the past by ourselves and others focusing on specific models for fibre orientation or diffusion tensor estimation (Xu et al. 2012; Hernández et al. 2013; Chang et al. 2014). In this paper we reformulate our previous proposed approach (Hernández et al. 2013) and provide a generic and model independent toolbox for model fitting using GPUs. The toolbox provides a flexible and friendly front-end for the user to specify a model, define constraints and any prior information on the model parameters and choose a non-linear optimisation routine, ranging from deterministic Gauss-Newton type approaches to stochastic Bayesian methods based on Markov Chain Monte Carlo (MCMC). It then automatically generates a GPU executable file that reflects all these options. This achieves flexibility in model fitting and accelerations of more than two orders of magnitude compared to CPU solutions.

To further explore the potential of GPUs for computational diffusion MRI, we present another parallel framework for white matter tractography and connectome estimation, a common dMRI application with completely different features and challenges compared to the voxel-wise biophysical modelling. We focus here on probabilistic tractography approaches, which for certain applications can be very time consuming (Behrens et al. 2007). For instance, the generation of a “dense” connectome (Sporns et al. 2005) from a single subject using high-resolution data from the HCP can take many days (see Table 1). In the case of tractography algorithms, the main challenge for a GPU parallel solution comes from the fact that the required data (set of voxels with distributions of fibre orientations) for propagating each streamline is not known in advance, as their paths are estimated dynamically on the fly. This makes the allocation of GPU resources difficult, and therefore, the a-priori assessment of the parallelisability of the application challenging. Moreover, the streamlines propagation is likely to be asynchronous as they may have imbalanced execution length, which induces thread divergence and causes performance degradation on GPUs. Furthermore, these methods have typically high memory requirements and include relatively heavy tasks, particularly for large datasets and whole-brain explorations, making the design of an efficient GPU-accelerated solution (which ideally comprises light and small tasks) even less straightforward. Preliminary GPU parallel tractography frameworks have been proposed in the past (Mittmann et al. 2008; Xu et al. 2012); however, our tractography parallel framework achieves accelerations of more than 200 times compared to CPU-based implementations and includes novel features that allow even more accurate anatomical constraints to be imposed, such as the inherent support of surface meshes (GIFTI files (Harwell et al. 2008)), and the possibility of generating dense connectomes.

In summary, we illustrate that, despite differences in parallelisability challenges, well-thought GPU-based designs that are carefully implemented can offer accelerations of more than two orders of magnitude, within the different contexts of tissue microstructure estimation, and tractography and connectome generation. The developed frameworks will be released upon publication within the FSL software library (Jenkinson et al. 2012) ^*^.

### THEORY

#### GPU architecture and CUDA

A GPU is a many-core device with its own memory space and connection to a host (CPU-based) system (see Figures 1 and 2 in the supplementary material for GPU architecture details) (Hernandez-Fernandez 2017). It is not a standalone platform but a co-processor typically connected to the host system via a Peripheral Component Interconnect express (PCIe) data bus (Budruk et al. 2004), a high performance I/O bus used to interconnect peripheral devices. A GPU contains thousands of computing cores, although the instruction set of these cores is much simpler than those on CPUs. It consists of a set of Single Instruction Multiple Threads (SIMT) *Streaming Multiprocessors* (SMs). The number of SMs on a GPU depends on the microarchitecture and the chip. A Streaming Multiprocessor (SM) is composed of many different units (NVIDIA 2015a), including processing units, memory resources and instructions dispatchers. The most important processing units in an SM are the *Streaming Processors* (SPs), which are responsible for most of the operations during the execution of a typical scientific application.

The GPU memory spaces are organised in a hierarchical way. The lower the level in the hierarchy the faster the memory access, but the smaller the storage space (NVIDIA 2015a; Cheng et al. 2014). At the highest level, we can find *global memory* (also called device memory). This is the largest storage space on a GPU device (up to 24 GB in Pascal NVIDIA architecture (NVIDIA 2016)), which is shared across all the SMs. Physically, it is not located on-chip (except for the new NVIDIA Pascal architecture (NVIDIA 2016)) and the access to this memory is expensive compared to the access to any other memory space. Global memory has high latencies (Andersch et al. 2014) and low bandwidth. The non-programmable *L2 cache* is located at an intermediate level. This coherent memory is located on-chip and is shared by all the SMs. Finally, at the lowest level we find several *cache memory* spaces (L1 cache, read-only/texture cache, Constant memory cache), *Shared memory and the register file*. All these memory spaces are inside an SM and they are shared by the threads allocated in it.

The CUDA programming model (Nickolls et al. 2008) allows programmers to use the GPU architecture for general parallel computing purposes. The key for GPU programming is multithreading using a data-level parallelism paradigm: thousands of threads execute the same instructions simultaneously over different data. A *kernel* is a function that is executed on the GPU by every thread (*single instruction multiple threads - SIMT*) launched from the host. CUDA is based on a hierarchy of abstraction layers. The whole set of threads that executes a kernel is called a *grid*. A grid is divided into groups of threads called thread-blocks or simply *blocks*. A block is mapped onto a single SM; therefore, the threads that belong to the same block will share the SM’s resources, including the processing units, registers, caches (excluding L2) and Shared memory. The threads within a block can communicate efficiently through Shared memory or the registers and they can be synchronised if necessary.

CUDA allows programmers to have control over the GPU resources and thus the possibility of designing specific parallel solutions for a certain application. However, programmers need to deeply understand the underlying parallel architecture and its components for taking advantage of the large amount of GPU’s computational power, making the development of efficient solutions challenging.

#### Considerations for GPU programming

The first consideration towards a parallel design for an application is to identify its most computationally expensive part and assess its parallelisability (Hernandez-Fernandez 2017). An application can be profiled for identifying these hotspots. The parallelisability of these hotspots will depend mainly on the number of independent parts which a task can be divided into. For completely exploiting the parallelism in modern GPUs, typically a minimum of 25,000 threads is required, and thus, not all types of applications are suitable for this architecture. Although GPU devices do not have so many cores (modern GPUs have around 4,000 cores), having more active threads than cores helps to hide latencies caused by memory accesses or floating point operations, and keeps the GPU cores always busy.

The threads within a CUDA block are grouped by the hardware into sets of 32 threads called *warps*. The warps are assigned simultaneously to an array of processing units and executed in a SIMT fashion, exploiting the data level parallelism. Thus, GPU parallel designs should follow an approach where all the subparts of the data are processed identically (i.e. same instructions), avoiding divergences. If a divergence occurs within a warp, for instance due to a branch or a loop, the different instructions will be sequentially computed by all the warp’s threads (deactivating some of them), and leading to performance degradation.

A CUDA programmer must decide the dimensions of the grid and blocks. This decision can have a significant impact on the performance obtained by an application executed on a GPU. The warp size (32 threads) imposes design constraints on the block and grid sizes, and there is as well a limitation on the maximum number of blocks simultaneously allocated. For maximum occupancy of resources (to allocate as many active threads as possible), the block size should be a multiple-of-32 number of threads, typically 128 or 258 threads. It is also important to keep in mind the communication and synchronisation requirements of the design. In general, we want to reduce communication and synchronisation as it consumes resources, but when necessary, we want this to happen within the same warp or alternatively within the same block, since CUDA offers relatively efficient and specific directives for these purposes at this level of the memory hierarchy.

The performance achieved by an application on a GPU does not only depend on its parallelisability or number of parts which a task can be divided into, but also on how efficiently these parts are executed on the device. CUDA programmers can control the memory spaces used for data storage. The use of low level spaces in the memory hierarchy should be prioritised, as the latencies are lower and the bandwidth higher. Moreover, the patterns of how pieces of data are requested by the threads within a warp can have a tremendous impact on the performance of an application, especially if these accesses are to global memory. During a memory access, each thread of a warp can request a different piece of data. A *memory segment* contains several pieces of data, but this is the minimum amount of data that can be transferred. Thus, if threads within a warp access to data allocated in different memory segments, all these segments will be transferred, even if most of the segment information is not used, leading to low memory access efficiency.

Once a part of the application has been accelerated, the next step is to identify a new bottleneck, i.e. the new part of the application that is consuming most of the computational time. The parallelised functions are normally implemented in different CUDA kernels, which actually can be processed in parallel on a GPU if they are independent.

## MATERIAL AND METHODS

In this paper we design and develop GPU frameworks for two different computational applications in diffusion MRI: a generic parallel framework for fitting biophysical/microstructure dMRI voxel-wise models, and a parallel framework for performing probabilistic tractography and generate connectomes. These two applications present very different challenges when it comes to the considerations outlined above. For the first application, we have designed a non-trivial two-level parallelisation solution that includes the collaboration of threads for fitting the model in a single voxel, and we develop a generic framework for easily designing, implementing and fitting MRI models. The second GPU framework, which performs probabilistic tractography presents completely different challenges, including the presence of thread divergence, the need for relatively heavy tasks, and high memory requirements. In our parallel design we propose solutions for these challenges.

### Biophysical Modelling on GPUs

#### Framework Description

Tissue microstructure estimation from dMRI is typically performed on a voxel-by-voxel basis, where a biophysical model is fitted. Excluding the DTI model, which can be easily and quickly fitted using linear least squares, most models are non-linear and numerical optimisation routines are required. Non-linear optimisation is typically computationally demanding and can be very time consuming, particularly since advanced multi-compartment models (Alexander et al. 2017) require larger than average datasets (multiple *b-values* or high angular resolution).

Given the large number of voxels and the relatively low memory requirements of these independent tasks, such an application is well-suited for implementation on GPUs. To take advantage of the inherent parallelisability of the problem and yet cover all possible dMRI models, we have developed a generic toolbox for designing and fitting nonlinear models using GPUs and CUDA. The toolbox, CUDA diffusion modelling toolbox (cuDIMOT), offers a friendly and flexible front-end for the users to implement new models without the need for them to write CUDA code or deal with a GPU design, as this is performed by the toolbox automatically (Figure 1). The user only specifies a model using a C-like language header. This model specification includes the model parameters, constraints and priors for these parameters, the model predicted signal function, and optionally the partial derivatives with respect to each parameter if they can be provided analytically (cuDIMOT offers the option for numerical differentiation). Once the model specification has been provided, the toolbox integrates this information with the parallel CUDA design of the corresponding fitting routines at compilation time, and it generates a GPU executable. The fitting routines include options for both deterministic (e.g. Gauss-Newton type) and stochastic (e.g. Bayesian inference using MCMC) optimisation.

**Figure 1.**
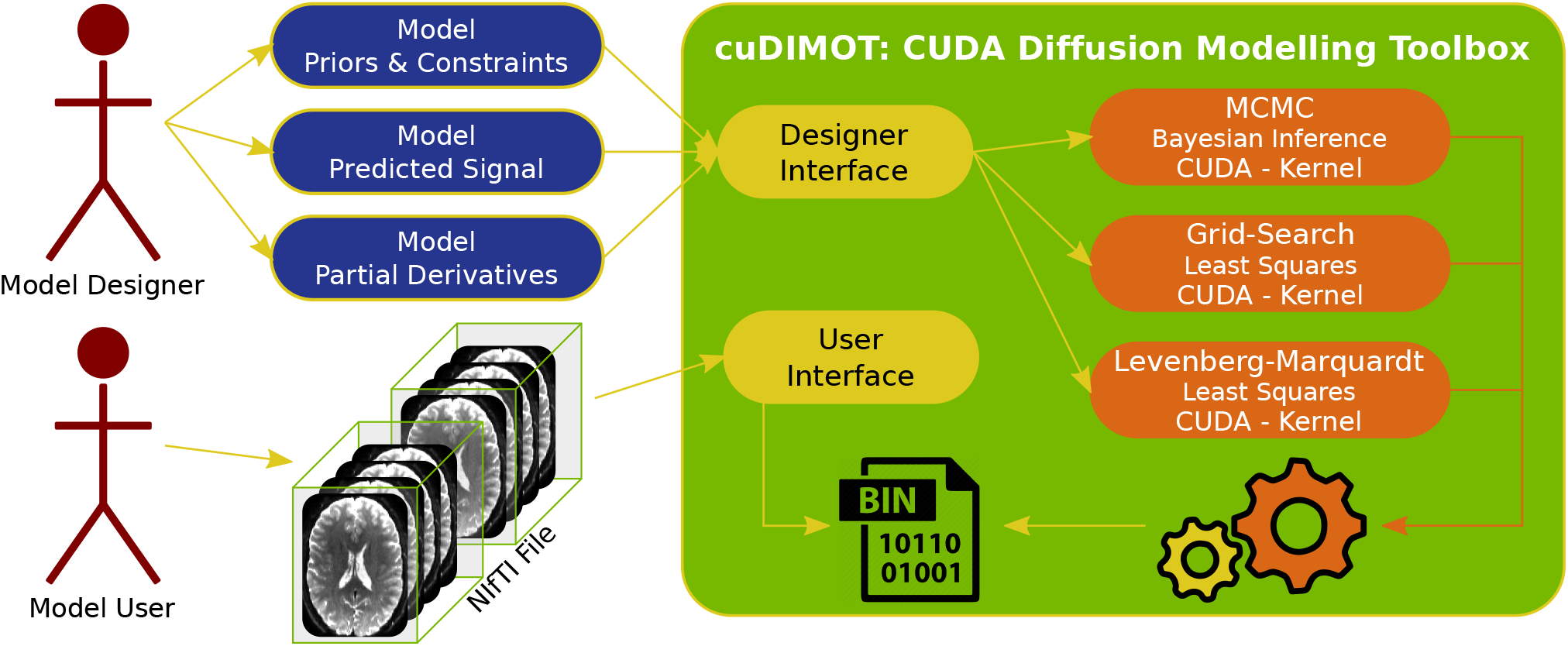
General design of CUDA Diffusion Modelling Toolbox (cuDIMOT). Two types of users interact with the toolbox through interfaces, a model designer and a model user. The model designer provides the model specification (parameters, priors, constraints, predicted signal and derivatives), whereas the model user interacts with the toolbox for fitting the model to a dataset. The toolbox provides CUDA kernels that implement several fitting routines. These kernels are combined with the model specification at compilation time for generating a GPU executable application.

#### Design considerations

An important factor to take into account in the design of the framework is its generic and model-independent aspect. In order to achieve an abstraction of the fitting routines, these are implemented in a generic way excluding functions that are model-dependent. The management of threads, definition of grid size and data distribution are also challenging aspects that cuDIMOT automates. The fitting routines, deterministic and stochastic, are implemented in different CUDA kernels. The toolbox implements the different kernels, deals with the arduous task of distributing the data among thousands of threads, uses the GPU memory spaces efficiently, and even distributes the computation among multiple GPUs if requested.

A number of facts complicate the optimal design of the toolbox: the specification of the model is known only at compilation time, and the dimensions of the dataset to be processed are not known until execution time. The first has driven a key aspect of the design, which is the integration of the model definition within the kernels. The CUDA kernels assume that there is a function *f*(***θ***), provided by the user. This function must return a scalar value with the model predicted signal given an array with the value of the model parameters ***θ***. For providing and defining *f*(***θ***) we decided to use external functions (implemented outside the optimisation routines). cuDIMOT is implemented using the CUDA C API, where the syntax for specifying the kernels and its functions is similar to C sequential functions. *f* (***θ***) is part of the CUDA kernels (integrated at compilation time), and thus, it can be implemented as a C function. The same logic is used for integrating partial derivatives of the cost function w.r.t. model parameters into the kernels. In this case, given an array with the value of the model parameters ***θ***, the function defined by the designer must return an array with several values, one per model parameter, which correspond to each partial derivative. Figure 2a shows an example of how a model is specified using the interface of the toolbox.

**Figure 2.**
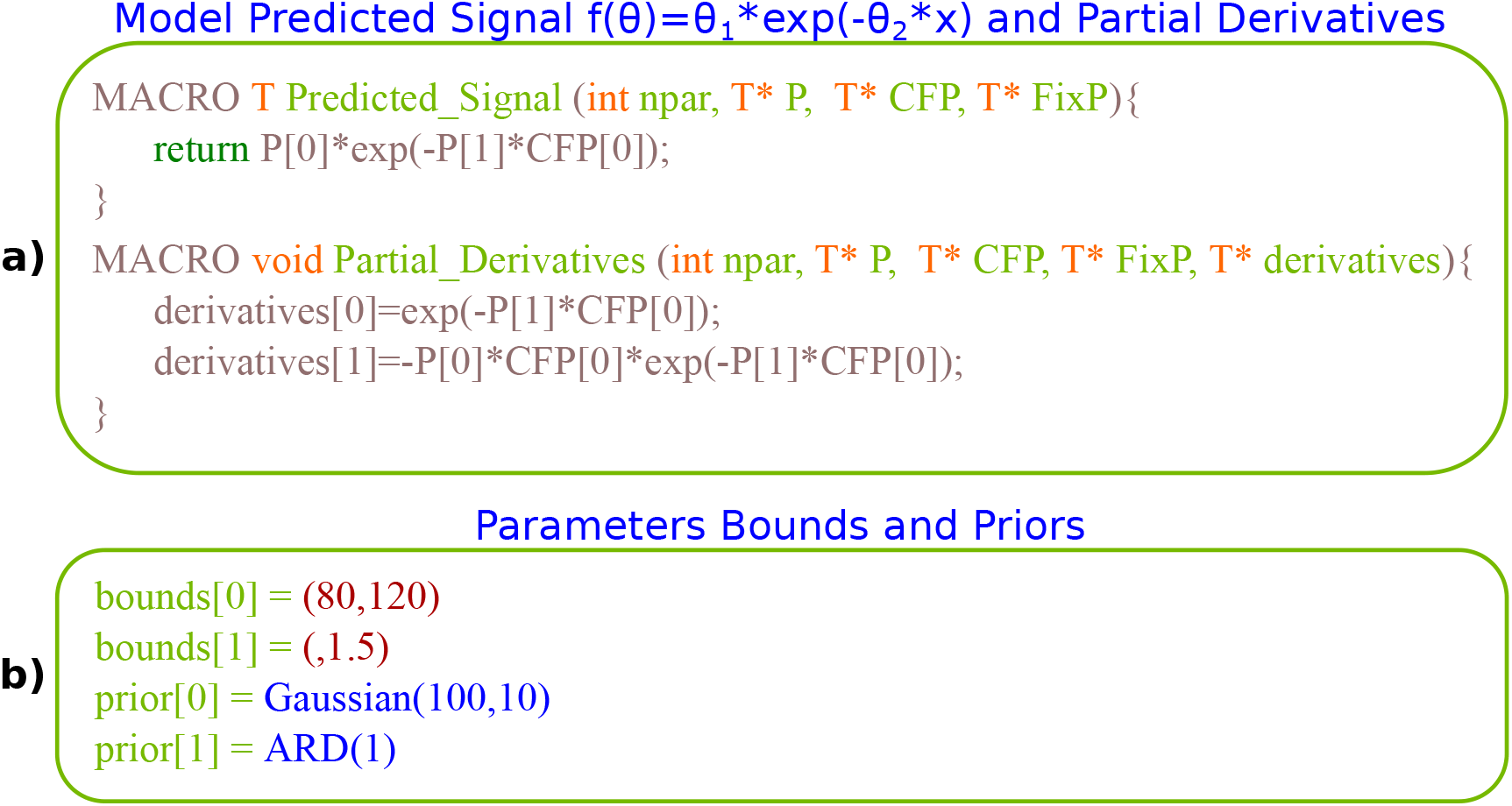
(a) Example of the specification of a simple model with 2 parameters in cuDIMOT. The model predicted signal returns a single value, and the function that defines the partial derivatives stores in an array the results for each parameter. The macro includes the necessary keywords for declaring a device function in CUDA, P contains the parameters Θ, and CFP and FixP contain the measurement points (CFP contains *x* in this specific function). (b) Prior information and constraints can be imposed on the model parameters.

Once the specification of the model has been performed, a GPU executable file is generated without the designer having to deal with any GPU considerations or code in CUDA. There is still one task that remains to be done, dealing with the distribution of the MRI data among the GPU threads. This cannot be done a-priori, as the dimensions of the dataset are only known at execution time. cuDIMOT implements a dynamic solution. When the kernels are executed, thousands of threads are created dynamically, as many as needed for processing the whole set of voxels. Internally, each thread has an ID, which is used to calculate, on the fly, the address of the data to process by each thread.

#### Model specification options

Models can have free parameters, which are estimated, and fixed parameters, which reflect measurement settings or features that are known. cuDIMOT allows such fixed parameters to be defined, and these may be common to all voxels or not (CFP or common fixed parameters and FixP or fixed parameters in Figure 2a respectively). For instance, in typical diffusion MRI, the diffusion-sensitising gradient strengths (b values) and associated directions (b vectors) would be CFPs, whereas for diffusion-weighted steady-state free precession (DW-SSFP) (McNab & Miller 2008), the flip angle (α) and repetition time (TR) would be CFPs, while the longitudinal and transverse relaxation times (T1 and T2) would be FixP, as they vary across voxels. Using a simple syntax, a list with all the information is passed to the toolbox through the designer interface. This information is parsed by cuDIMOT and used at execution time, when the model user must provide maps with these parameters. This generic interface allows the users to combine data from dMRI with data from other modalities, such as relaxometry (Deoni 2010), and develop more complex models (Foxley et al. 2016; Foxley et al. 2015), or, even use cuDIMOT in different modalities where nonlinear optimisation is required.

In terms of optimisation routines, cuDIMOT offers a number of deterministic and stochastic model-fitting approaches, including greedy optimisation using Grid-Search, non-linear least-squares optimisation using Levenberg-Marquardt (LM) and Bayesian inference using Markov Chain Monte Carlo (MCMC). Prior information or constraints on the parameters of a model can be integrated into the fitting process using the toolbox interfaces, where a simple syntax is used for enumerating the type and the value of the priors (see Figure 2b). Upper and lower limits or bounds can be defined for any parameter (transformations are implemented internally and described in the supplementary material), and priors can be any of the following types:

- A Normal distribution. The mean and the standard deviation of the distribution must be specified.
- A Gamma distribution. The shape and the scale of the distribution must be specified.
- Shrinkage prior or Automatic Relevance Determination (ARD) (MacKay 1995). The factor or weight of this prior must be provided in the specification.
- Uniform distribution within an interval and uniform on a sphere (for parameters that describe an orientation).

For stochastic optimisation, a choice can be also made on the noise distribution and type of the likelihood function (Gaussian or Rician).

Internally, a *C++* parser was also implemented to read and store the list of these options, which are checked at runtime during the fitting process. It is also possible to define constraints that depend on the relation of two or more parameters (for instance, a < b). These constraints can be specified using external functions and the C syntax, similar to the specification of the model predicted signal function.

Because high-dimensional models are difficult to fit without a good initialisation, the toolbox offers an option for cascaded fitting, where a simpler model is fitted first and the estimated parameters are used to initialise the parameters of a more complex model. 3D volumes can be used for specifying the initialisation value of the model parameters in every voxel.

Once a model has been defined and an executable file created, a user still has flexibility in controlling a number of fitting options, including:

- Choosing fitting routines: Grid-Search, Levenberg-Marquardt or MCMC. A combination of them is possible, using the output of one to initialize the other.
- Selecting number of iterations in Levenberg-Marquardt and MCMC (burn-in, total, sample thinning interval).
- Using Gaussian or Rician noise modelling in MCMC.
- Choosing model parameters to be kept fixed during the fitting process.
- Choosing model selection criteria to be generated, such as BIC and AIC.

#### Challenges and Parallel design

As described above, a straightforward approach towards a GPU parallel design is motivated by the independent nature of the fitting process across voxels. A naive parallel design can assign each voxel fitting process to a different CUDA thread. However, such an approach has considerable limitations. The within voxel computations lead to many operations and relatively heavy-weight threads. This means that parallelism will be low as fewer threads can be allocated simultaneously on the GPU. *Uncoalesced* memory accesses is another concern. Voxel-wise data is too large to fit into registers or Shared memory, and thus has to be stored in global memory. In this design, threads within a warp access data from different voxels, which is located in different memory segments (segments store diffusion measurements from the same voxel) and this causes many memory transfers. All these factors considerably reduce performance.

We found that a second level of parallelisation (see Figure 3) increases massively GPU occupancy and therefore performance. Similar to our previous work (Hernández et al. 2013), cuDIMOT parallelises the computation of the most expensive within-voxel tasks, which is the cost function calculation in the case of deterministic optimisation, or equivalently, the likelihood calculation in the case of stochastic optimisation. Specifically, in the first level of parallelisation, the fitting process of a group of *B* voxels is assigned to a CUDA block. Subsequently, in the second level of parallelisation the fitting process of each voxel is assigned to a warp. Threads within a warp collaborate in the following tasks:

- Compute the model predicted signal and residuals across the different measurement points in MCMC, Levenberg-Marquardt, and Grid-Search
- Calculate partial derivatives with respect to the model parameters across the different measurement points in Levenberg-Marquardt
- LU decomposition in Levenberg-Marquardt. The LU decomposition is used for solving the system:

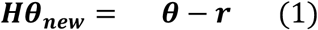

for calculating the proposed parameters ***θ_new_*** in every iteration of the Levenberg-Marquardt algorithm, where ***r*** is the gradient and **H** the approximated Hessian matrix. Using LU decomposition in the matrix ***H***:

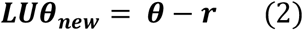

In cuDIMOT, we solve this system using *P +* 1 threads, where *P* is the number of columns of ***H*** (number of model parameters). Figure 4 depicts the parallelisation of the method, which uses Gaussian elimination with pivoting and solves the system as:

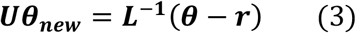

**Figure 3.**
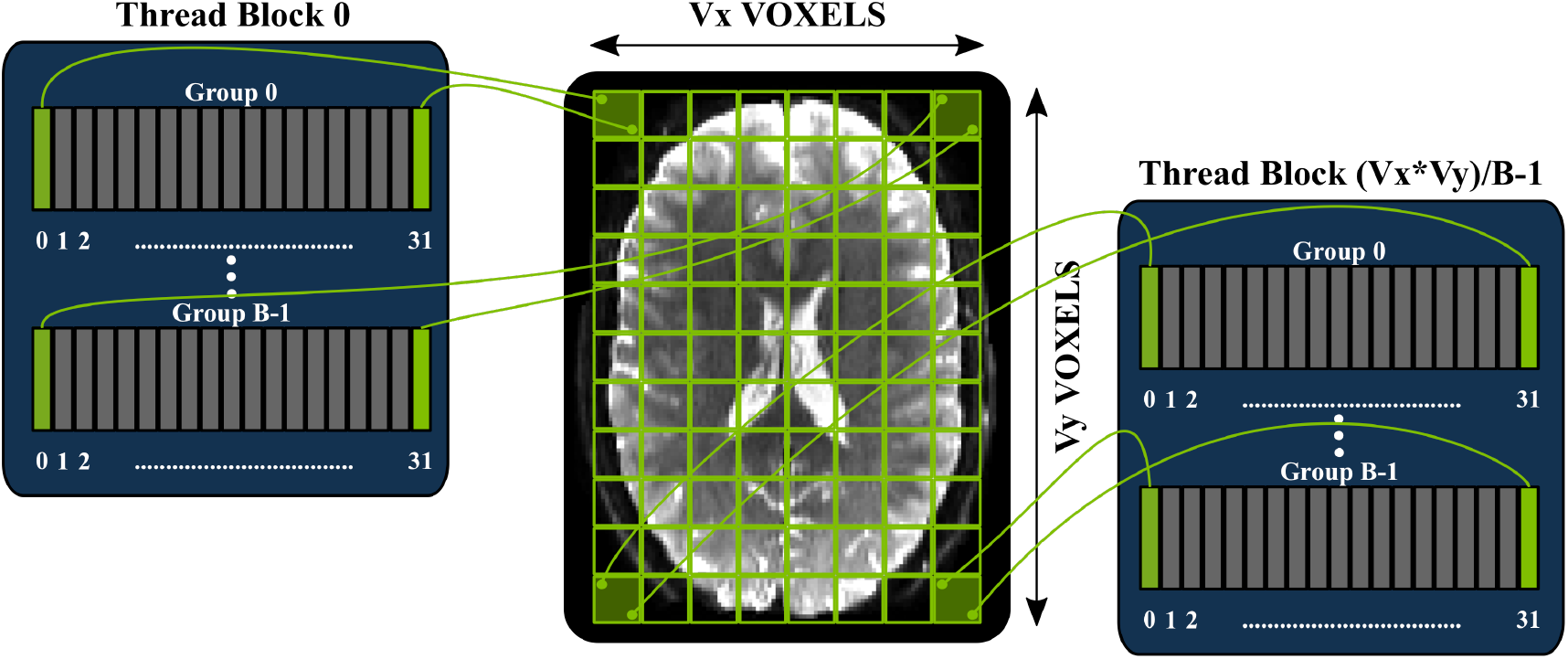
Parallel design of cuDIMOT for fitting dMRI models on a GPU. The V voxels of a dataset are divided into groups of B voxels (voxels per block), and the fitting process of each of these groups is assigned to different CUDA blocks. Inside a block, a warp (32 threads) collaborate for within-voxel computations.

**Figure 4.**
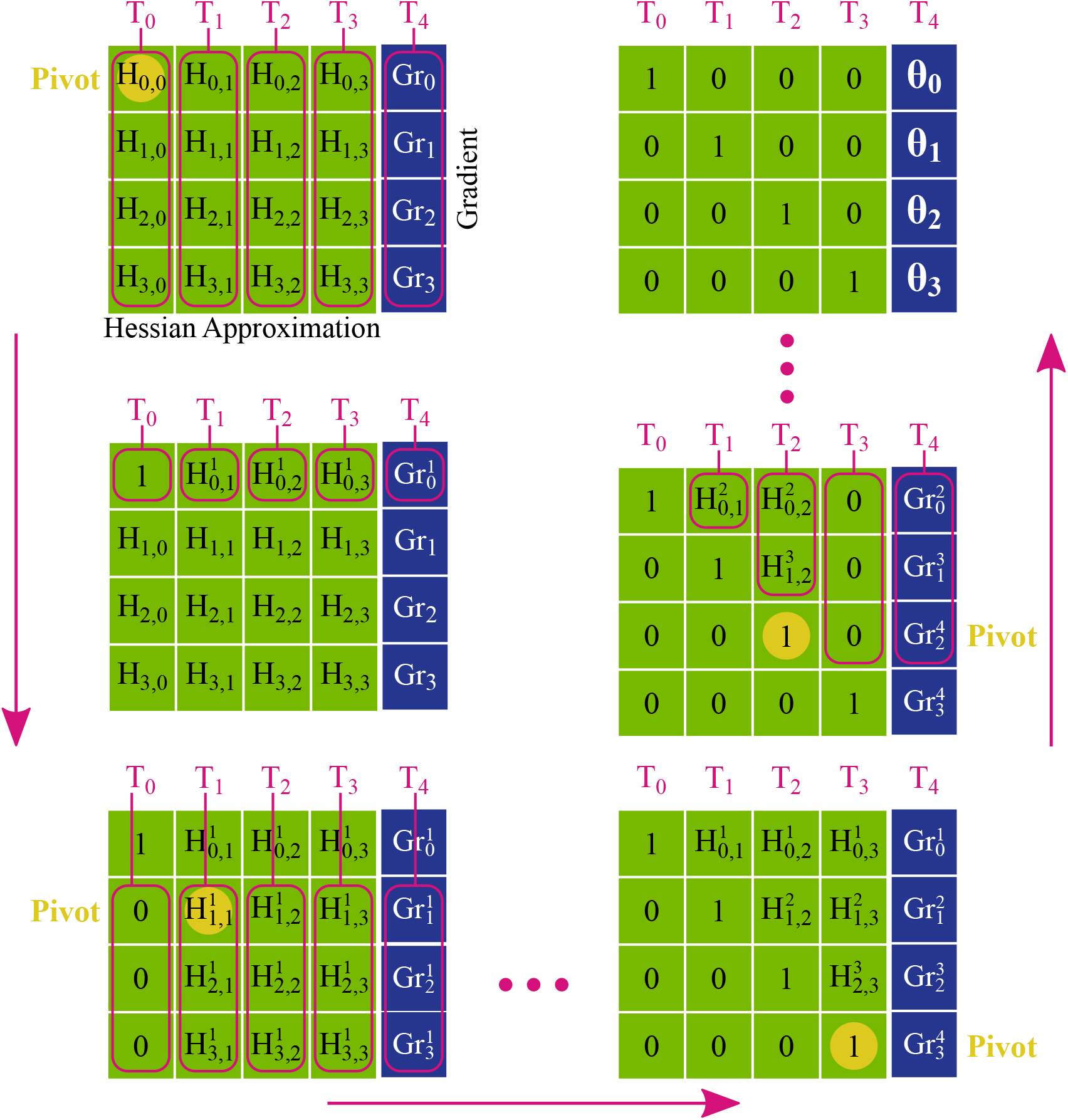
Parallelisation of the LU decomposition across the threads within a warp. The LU decomposition is computed using Gaussian elimination with pivoting. The method produces ones at the diagonal elements of the matrix and zeros above and below the diagonal. The results are produced at the rightmost column. Each thread T computes in parallel the elements of a column.

Firstly, *P* forward steps are made for calculating ***U***, producing ones at the diagonal of the matrix and zeros below the diagonal. At each step a diagonal element is set as the pivot and each thread computes the elements of a different column. CUDA shuffle instructions (NVIDIA 2015a) are used for broadcasting the pivot. CUDA shuffle instructions allow threads to read the registers of other threads belonging to the same warp. This communication mechanism is faster than Shared memory and it does not require synchronisation, as it is implicit within the threads of a warp. Secondly, *P* backward steps are made for producing zeros above the diagonal. Again, each thread computes the elements of a different column. The method produces the results ***θ_new_*** at the rightmost column.

It is easy to see that the threads compute different number of elements at each step of the method, i.e. the workload is not balanced across threads. Also, the number of columns of the matrix is most often lower than the number of threads of a warp, and thus there are idle threads. Nevertheless, this implementation is more efficient than solving the system with only one CUDA thread.

The threads that cooperate for computing the fitting process in a voxel need to communicate with each other frequently, for instance for summing the residuals and compute the cost function. We use shuffle instructions for performing this communication.

This two-level design addresses most of the previous limitations:

- It does not involve heavy tasks. Now threads are very light and they consume less computational resources.
- The most expensive operations are now processed by a group of threads instead of by a single thread, exposing more parallelism.
- Memory accesses are coalesced. When threads within a warp access to the diffusion measurements (from the same voxel), these are in consecutive memory locations and thus the efficiency of memory accesses increases significantly.

A higher-level of parallelism can be further used to enhance even more the performance in the framework, using very large groups of voxels and a multi-GPU system. We can divide a single dataset into groups of voxels and assign each group to a different GPU. The different GPUs do not need to communicate because the groups of voxels are completely independent, apart from the final step of writing the results. Typically, this can also be applied to a CPU multi-core processor or to a cluster of processing nodes.

#### Exploring microstructure diffusion MRI models with cuDIMOT

We used cuDIMOT for implementing a number of diffusion MRI models and assess the validity of the results. We have implemented the Neurite Orientation Dispersion and Density Imaging (NODDI) model, using Watson (Zhang et al. 2012) and Bingham (Tariq et al. 2016) distributions for characterising orientation dispersion.

We implemented NODDI-Watson with cuDIMOT using the designer interface. This model assumes the signal comes from three different compartments: isotropic compartment, intra-cellular compartment and extra-cellular compartment. The model has five free parameters: the fraction of the isotropic compartment *f_iso_*, the fraction of the intra-cellular compartment relative to the aggregate fraction of the intra-cellular and extra-cellular compartments *f_intra_*, the concentration of fibre orientations *κ* (the lower this value the higher the dispersion), and two angles for defining the mean principal fibre orientation *θ* and ϕ. The concentration parameter *κ* can be transformed and expressed as the orientation dispersion index *OD ϵ* [0,1]:

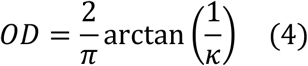

We implemented the model predicted signal of NODDI-Watson as in (Zhang et al. 2011), providing analytically the derivatives for *f_iso_*, *f_intra_*. We used numerical differentiation to evaluate the partial derivatives of the rest of the parameters. We used numerical approximations (e.g. for the Dawson’s integral) as in (Press et al. 1987), and we performed the same cascaded steps as the Matlab NODDI toolbox (Microstructure Imaging Group-University College London n.d.). First, we fit the diffusion tensor model for obtaining the mean principal fibre orientation (*θ* and ϕ). Second, we run a Grid-Search algorithm testing different combination of values for the parameters *f_iso_*, *f_intra_* and *κ*. Third, we run Levenberg-Marquardt algorithm fitting only *f_iso_*, *f_intra_*, and fixing the rest of parameters. Finally, we run Levenberg-Marquardt, fitting all the model parameters. The only difference is that the Matlab toolbox uses an active set optimisation algorithm (Gill et al. 1984) instead of Levenberg-Marquardt.

The NODDI-Bingham model assumes the same compartments as NODDI-Watson. However, this model can characterise anisotropic dispersion, and thus it has two concentration parameters *κ*_1_, *κ*_2_, and an extra angle *ψ* which is orthogonal to the mean orientation of the fibres, and it encodes a rotation of the main dispersion direction. When *κ*_1_ = *κ*_2_ the dispersion is isotropic, and when *κ*_1_ > *κ*_2_ anisotropic dispersion occurs. In this case the orientation dispersion index *OD* is defined as:

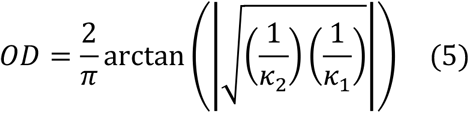

and a index *DA ϵ* [0,1] reflecting the factor of anisotropic dispersion can be defined as:

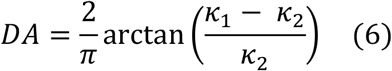

We implemented NODDI-Bingham using cuDIMOT in a similar manner as the previous model (only providing the analytic derivatives for the *f_iso_* and *f_intra_* parameters). For implementing the confluent hypergeometric function **1*𝓕***_**1**_ of a matrix argument included in the predicted signal of the intra-cellular compartment of the model, we use the approximation described in (Kume & Wood 2005). We use the same optimisation steps as in the NODDI-Watson: diffusion tensor fitting, Grid-Search, and Levenberg-Marquardt twice.

### Probabilistic Tractography and Connectomes on GPUs

Contrary to voxel-wise model fitting, white-matter tractography, and particularly whole-brain connectome generation, are not inherently suited for GPU parallelisation. Very high memory requirements, uncoalesced memory accesses and threads divergence (irregular behaviour of threads in terms of accessed memory locations and life duration) are some of the major factors that make a GPU parallel design of such an application challenging. Nevertheless, we present a framework that parallelises the propagation of multiple streamlines for probabilistic tractography and overcomes the aforementioned issues using an overlapping pipeline-design.

#### Framework Description

We design a parallel design and develop a GPU-based framework for performing probabilistic tractography. Our application includes the common tractography functionality, including for instance options to set:

- The number of streamlines propagated from each seed point, i.e., the number of samples.
- Streamline termination criteria (maximum number of steps, curvature threshold, anisotropy threshold, tract loop detection)
- A number of numerical integration approaches, including Euler’s method and 2^nd^ order Runge-Kutta method (Basser et al. 2000), with a subsequent choice of step length.
- Propagation criteria and anatomical constraint rules (seed, waypoint, termination, stopping, and target masks) (Smith et al. 2012).
- Ability to accept tracking protocols in either diffusion or structural/standard space.

Connectome generation is also inherently supported (Sporns et al. 2005) and three options are available (see Figure 5) (Li et al. 2012; Donahue et al. 2016):

- Connectivity matrix between all the seed points and all the other seed points. A typical use is for studying the connectivity from all grey matter to all grey matter (Glasser et al. 2013).
- Connectivity matrix between all the points of two different masks, which are independent of the seed points mask. A typical use is for studying the connectivity between grey matter regions, when seeding from all white matter.
- Connectivity matrix between all the seed points and all the points specified in a different mask. A typical example is to use the whole brain as a target mask for studying the connectivity profile of the grey matter in a specific seed, and use it for connectivity-based classification (Johansen-Berg et al. 2004).

**Figure 5.**
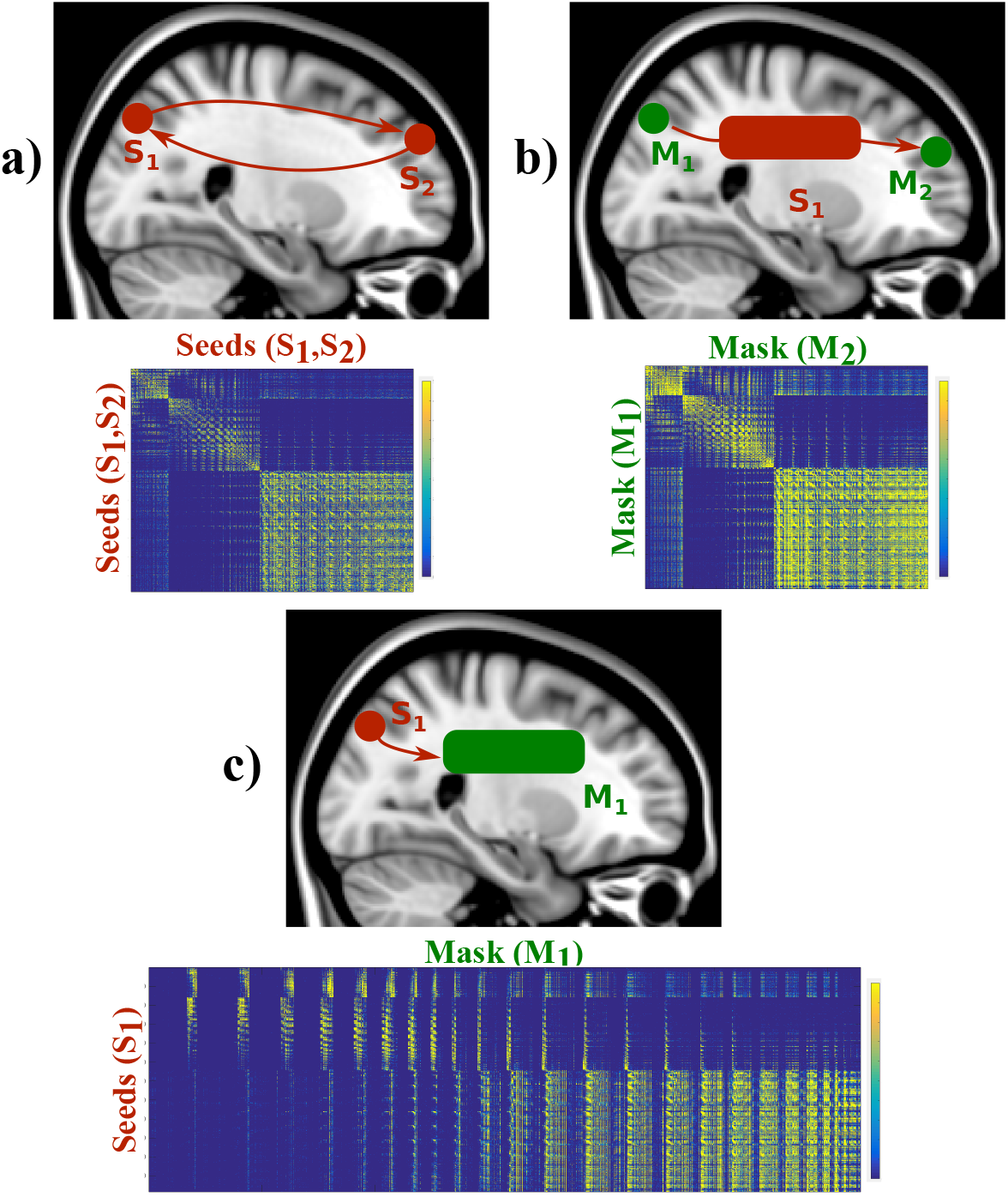
Connectivity matrices modes offered by the GPU-accelerated tractography framework. The framework can generate connectivity matrices from a) all seed to all seed points, b) all points in a mask to all points in another mask seeding from an independent region, or c) all seed points to all points in a different mask.

We have also included an extra feature, which is not typically found in tractography toolboxes, but can be important in defining anatomically accurate constraints. We included in our parallel framework the possibility of using surfaces, as well as volumes, for defining seed and regions of interest (ROIs). We implement support for the GIFTI format (Harwell et al. 2008), according to which surfaces are defined by meshes of triangles. Three spatial coordinates define each triangle vertex in a 3D space. Surface vertices can be used for defining seed points, and mesh triangles can be used for defining stopping/constraint masks. If the latter, a streamline needs to be checked upon crossing the surface meshes. We implement the method described in (O’Rouke 1998) (ray-plane intersection) for checking if the segments of a streamline intersects a triangle (details of the method are presented in the supplementary material).

#### Challenges and Parallel design

Path propagation is completely independent across streamlines, and thus it can be in principle parallelised. To reflect that, we can create as many CUDA threads as the required number of streamlines (in fact twice the number of streamlines, as we propagate from each seed point towards both directions indicated by the local fibre orientation). Thus, for *D* seed points and *F* streamlines per seed, we have *2 x D x F* threads in total (see Figure 6). Nevertheless, there are complexities that make such a design challenging and considerably reduce efficiency if not addressed, as we explain below. These include heavy tasks, thread divergence and memory allocation challenges.

**Figure 6:**
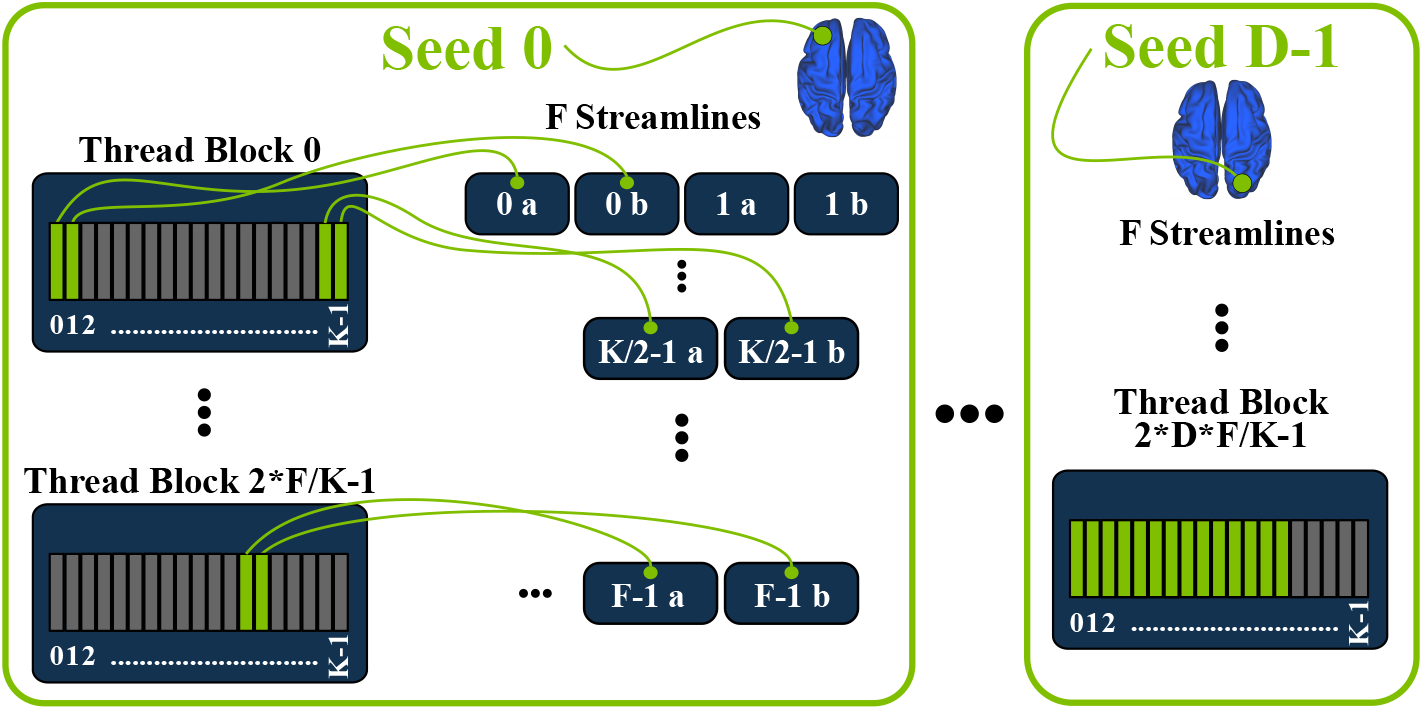
GPU parallel design of the streamline probabilistic tractography framework. For each of the D seeds and for each of the F streamlines per seed, we create two CUDA threads (a and b), which are distributed amongst blocks of K threads.

A first consideration is the high complexity of some of the routines included for implementing the offered functionality. For instance, the use of surfaces involves the execution of an intersection detection algorithm, while the streamline propagation includes interpolation and space transformation routines. Furthermore, checking anatomical constraints during propagation increases the complexity of the algorithm, and induces a significant number of conditional branches. Having a single CUDA kernel for performing all these tasks leads to substantially heavy threads, which consume a lot of computational resources and consequently cause low occupancy in a SM.

To solve this issue, we split the application into multiple tasks, each of which is implemented in a different CUDA kernel. Specifically, we have designed the following:

- A kernel that propagates the streamlines and checks the basic termination criteria, including curvature and anisotropy thresholds, loop detection and out of brain detection.
- Several kernels, for checking the anatomical constraints masks. We have a different CUDA kernel for each type of anatomical constraint, i.e., a kernel for checking the stopping masks, a kernel for checking the exclusion masks, a kernel for checking the inclusion masks and a kernel for checking the target masks.
- A kernel for updating the path distribution (visitation count) map. It uses the streamlines’ coordinates for updating each visited voxel in a unique and shared volume of voxels located in the GPU global memory.
- A kernel for generating connectomes. This optional kernel checks if any of the elements of the volumes and surfaces of the connectivity matrix has been visited. For “dense” connectomes connectivity matrices can be very large (~30 GB in the case of using HCP grayordinates (Glasser et al. 2013)) and they cannot be allocated onto the GPU memory. Thus, each thread of this kernel stores in a vector only the IDs of the visited matrix elements. The vectors are transferred from the GPU to the host, which finally executes a routine for updating a matrix allocated in the host memory.

In order to execute all these kernels, the pipeline depicted in Figure 7a is used, where each kernel runs serially one after the other. The locations (coordinates) of the streamlines, obtained in the propagation kernel, need to be stored for use in the other kernels, which increases the memory requirements (3 floats per coordinate). However, this is not particularly problematic compared to the heavy task that would have to be performed using a single kernel. With the proposed multi-kernel design we achieve much lighter CUDA threads, relieving pressure on the number of used registers per thread.

**Figure 7.**
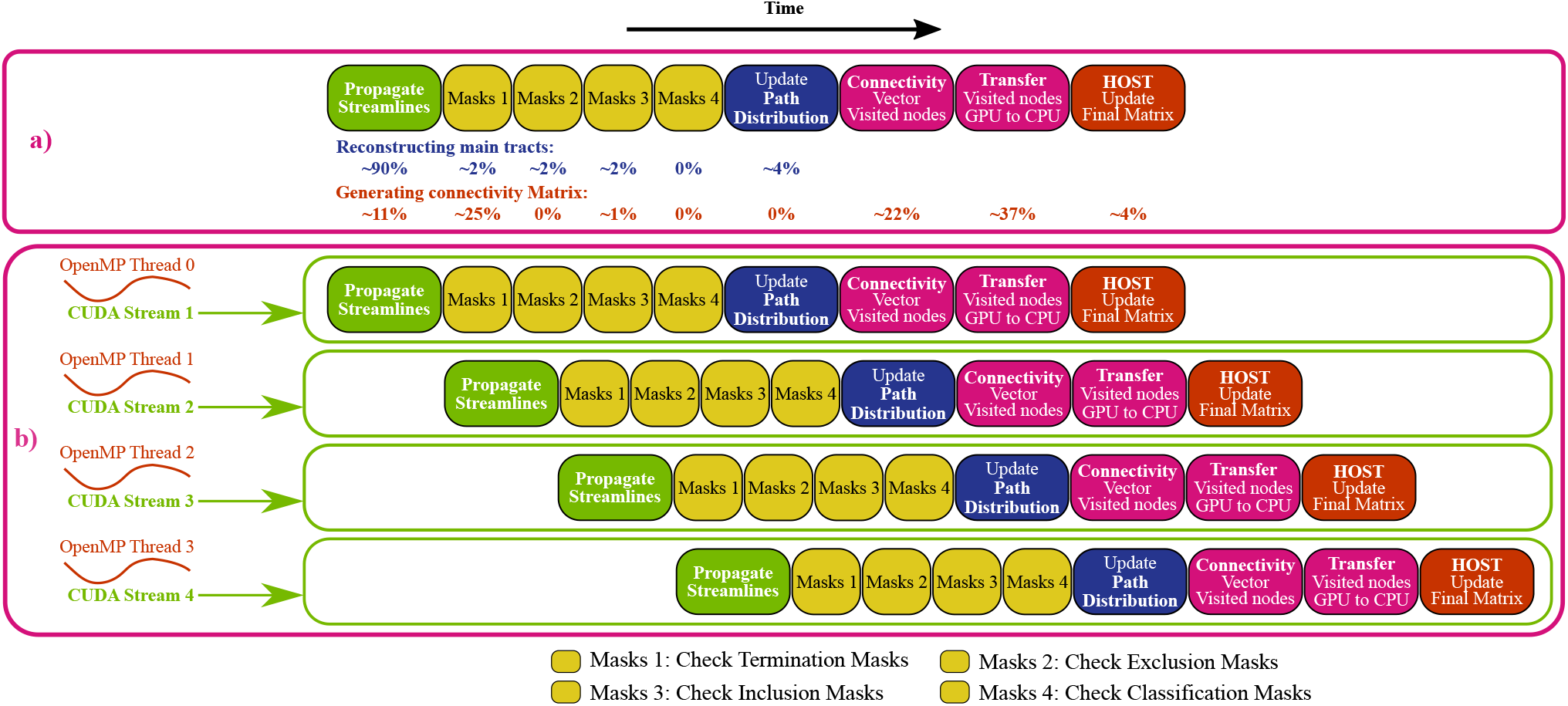
a) Pipeline of the GPU-accelerated probabilistic tractography framework. The routines that implement the different functionality of the application are split into different CUDA kernels. The connectivity matrix is stored in the host memory, and the GPU threads only store the visited nodes of the matrix (in a vector). The host needs to execute a final process for generating and updating the connectivity matrix. The figure shows the percentage of time spent on each kernel when typical tracts are reconstructed, and when a grey matter to grey matter dense connectome is generated. In both cases, no classification mask was used, and for generating the connectome, no exclusion mask was employed and the path distribution was not generated, thus, 0% of time was spent on these kernels. b) Overlap of several pipelines in the GPU-accelerated tractography framework using several CUDA streams and OpenMP threads.

Another main challenge is related to memory requirements. It may be impossible to use GPUs if the memory demands exceed the available device memory, and typically, this is true across all levels in the GPU memory hierarchy. Tractography algorithms need a distribution of fibre orientations for sampling. As we cannot predict the streamline track locations in advance, the fibre orientation distributions of all voxels need to be allocated in memory. The amount of required memory depends on the size (spatial dimensions) of the dataset and for probabilistic tracking on the number of samples in the orientation distributions. For instance, the required memory for simply allocating the samples of a Human Connectome Project dataset (Van Essen & Ugurbil 2012; Van Essen et al. 2012; Sotiropoulos et al. 2013) is approximately 1.5 GB. Moreover, the 3D streamline coordinates need to be stored, but the number of steps that a streamline will take cannot be predicted in advance. We therefore need to allocate enough memory for storing the maximum possible number of coordinates for each streamline. Additionally, volumes and/or surfaces may be used for defining seeds, anatomical constraints and connectome matrix elements, and all of them need also to be allocated in memory. Our strategy for overcoming this issue was to allocate all the required memory in the GPU without considering the streamline coordinates, and then calculate the maximum number of streamlines that can be computed in parallel given the amount of memory left. If all the requested streamlines cannot be computed in parallel (which is the most typical scenario), the application iterates over different streamline sets.

All the memory allocation is initially performed in global memory, which is suboptimal and causes high latencies every time the threads access to it. For optimizing the memory transfers between host and device, we allocate the data in the host as *pinned* memory, which forces the data to be page-locked, allowing faster transfers to and from the GPU. It is more expensive to allocate this memory, but once allocated the transfers become faster in consecutive iterations for processing the next streamline sets. In addition, to mitigate latencies when accessing global memory, we use as much as possible the most efficient memory spaces, such as Shared memory. We store the most commonly accessed data in Shared memory: the parameters of the streamlines that are needed for propagation (current coordinates, and direction in which propagation will continue). Similarly, we store space transformation information in the efficient read-only constant memory.

Another challenge that limits performance is thread divergence. The streamlines may be initialised at different seed points and they can propagate to different regions and for different lengths. This causes accesses to different memory locations when sampling, i.e., uncoalesced memory accesses. Furthermore, streamlines may terminate at different time points. This causes asynchronous thread termination and a possible waste of computational resources, as some threads finish their execution before others and stay idle, wasting GPU resources. This situation persists until all the threads of the same block finish their execution and the GPU scheduler switches the block with another block. For this reason, we set the block size K to a small size of 64 threads. Although with this configuration full SM occupancy is not achieved (because there is a limit in the number of blocks per SM), there can be fewer divergences than when having larger blocks. Moreover, the GPU can employ the unused resources by overlapping other tasks/kernels in parallel.

We indeed take advantage of our pipelined design and use *CUDA streams* to overlap the computation of different sets of streamlines. CUDA streams are queue instances for managing the order of execution of different tasks (NVIDIA 2015a) and offer the possibility of running concurrently several tasks (kernels execution and/or CPU-GPU memory transfers) on the same GPU. Our framework divides the streamlines into a few sets, and uses a number of OpenMP (Chapman et al. 2008) threads to execute the pipeline of several streamline sets on different CUDA streams concurrently^†^ (see Figure 7b).

### Diffusion–weighted MRI data

Both GPU designs presented here have been used so far to process hundreds of datasets. Here, we illustrate performance gains on various exemplar data with both high and low spatial resolutions.

For testing the diffusion modelling framework (cuDIMOT) we used data from UK Biobank (Miller et al. 2016). Diffusion-weighting data were acquired using an EPI-based spin-echo pulse sequence in a 3T Siemens Skyra system. A voxel size of 2.0×2.0×2.0 mm^3^ was used (TR = 3.6 s, TE = 92 ms, 32-channel coil, 6/8 partial Fourier) and 72 slices were acquired. Diffusion weighting was applied in M = 100 evenly spaced directions, with 5 directions b = 0 s/mm^2^, 50 directions b = 1,000 s/mm^2^ and 50 directions b = 2,000 s/mm^2^. A multiband factor of 3 was employed (Moeller et al. 2010; Setsompop et al. 2012). A T1 structural image (1 mm isotropic) of the same subject was used for creating a white & grey matter mask, which was non-linearly registered to the space of the diffusion dataset (Andersson et al. 2007), and applied to the map of the estimated parameters before showing the results included here. For creating this mask, a brain extraction tool (Smith 2002) and a tissue segmentation tool (Zhang et al. 2001) were used.

For testing the GPU probabilistic tractography framework, data were acquired on a 3T Siemens Magnetom Prisma using HCP-style acquisitions (Sotiropoulos et al. 2013). Diffusion-weighting was introduced using single-shot EPI, using an in-plane resolution of 1.35×1.35 mm2 and 1.35 mm slice thickness (TR=5.59 s, TE=94.6 ms, 32-channel coil, 6/8 partial Fourier). 134 slices were acquired in total and diffusion weighting was applied in M = 291 evenly spaced directions, with 21 directions b = 0 s/mm2, 90 directions b = 1,000 s/mm2, 90 directions b = 2,000 s/mm2 and 90 directions b = 3,000 s/mm2.

### Hardware Features

We used an Intel host system with NVIDIA GPUs and a large cluster of Intel processors for testing our parallel designs. The former system is comprised of 2 Intel Xeon E5-2680 v3 2.50 GHz processors, each with 12 cores (24 CPU cores in total and 48 threads), and with 384 GB (24 x 16 GB) RDIMM memory. The system has 2 NVIDIA K80 accelerators (Error Correcting Codes ECC enabled), each connected to a different processor via PCI express v3. Each NVIDIA K80 accelerator has 2 GPUs, thus the system has in total 4 GPUs. The second cluster has tens of CPU nodes, and each node is comprised of 2 Intel Xeon E5-2680 v3 2.50 GHz processors (the same processors as the other system), each with 12 cores (24 CPU cores per node), and 320 GB RDIMM memory. Major features of the NVIDIA K80 accelerators and the Intel processors are summarized in Table 2 (NVIDIA 2014a; NVIDIA 2015b).

**Table 2.** Major features of NVIDIA Tesla K80 accelerator (2 GPUs) & Intel Xeon E5-2680 v3 processor (10 cores).

| Element | Feature | NVIDIA Tesla K80 accelerator |
| --- | --- | --- |
| Processing Units | Number of GPUs per accelerator | 2 (Tesla GK210) |
|  | Number of SMs per GPU | 13 |
|  | Single precision cores per SM | 192 |
|  | Double precision cores per SM | 64 |
|  | Special Function Units per SM | 32 |
|  | Load/Store Units per SM | 32 |
|  | Texture Units per SM | 16 |
|  | Core clock | 560 MHz |
| Memory | DRAM - Main memory per GPU | 12 GB |
|  | L2 Cache per GPU | 1,536 KB |
|  | Shared memory per SM | 80/96/112 KB |
|  | L1 cache per SM | 16/32/48 KB |
|  | Read Only Cache per SM | 48 KB |
|  | Constant memory cache per SM | 10 KB |
|  | Register File per SM | 512 KB |
|  | Memory clock | 2,500 MHz |
|  | Bandwidth per GPU | 240 GB/s |
| Schedulers/Dispatchers | Warp Schedulers per SM | 4 |
|  | Instruction dispatch units | 8 |
| Processing Power | Peak single precision | 2 x 4.365 TFLOPS |
|  | Peak double precision | 2 x 1.45 TFLOPS |
| Power Consumption |  | 300 watts |
| Price (January 2017) |  | £4,960 |
| Element | Feature | Intel Xeon E5-2660 v3 |
| Processing Units | Number of Cores (per processor) | 10 |
|  | Clock frequency | 2.60 GHz |
| Memory | DRAM - Main Memory (system) | 320 GB |
|  | L1 Cache (per processor) | 25 MB |
| Processing Power per processor | Peak single precision | 832 GFlops |
|  | Peak double precision | 416 GFlops |
| Processing Power per single core | Peak single precision | 83.2 GFlops |
|  | Peak double precision | 41.6 GFlops |
| Power Consumption |  | 105 watts |
| Price (January 2017) |  | £1,500 |

These systems run Centos 6.8 Linux. We compiled our code using CUDA 7.5 (V.7.5.17) and gcc 4.4.7 compilers. Experiments were performed on the system, with the sequential version of the algorithm running on a single CPU core (and running a single thread) and the parallel versions on a single GPU. Multi-GPU and multi-core experiments were also performed using all the available CPU cores and all the available GPUs (4) in the system.

## RESULTS

### Tissue microstructure modelling with GPUs

We fit the NODDI-Watson model to a UK Biobank dataset using three approaches, the NODDI Matlab toolbox (Microstructure Imaging Group - University College London n.d.), AMICO (Daducci et al. 2015) and cuDIMOT. The NODDI toolbox can parallelise the fitting process distributing groups of voxels among several CPU threads. AMICO reformulates the problem as a linear system via convex optimisation and accelerates computations by performing discrete searches in the multidimensional space of the problem. Figure 8a shows maps with the estimated parameters from each approach and respective execution times. Both cuDIMOT and AMICO achieved accelerations of more than two orders of magnitude compared to NODDI toolbox (cuDIMOT 352x and AMICO 160x) using a single NVIDIA K80 GPU and a single CPU core respectively. cuDIMOT was 2.2 times faster than AMICO.

**Figure 8.**
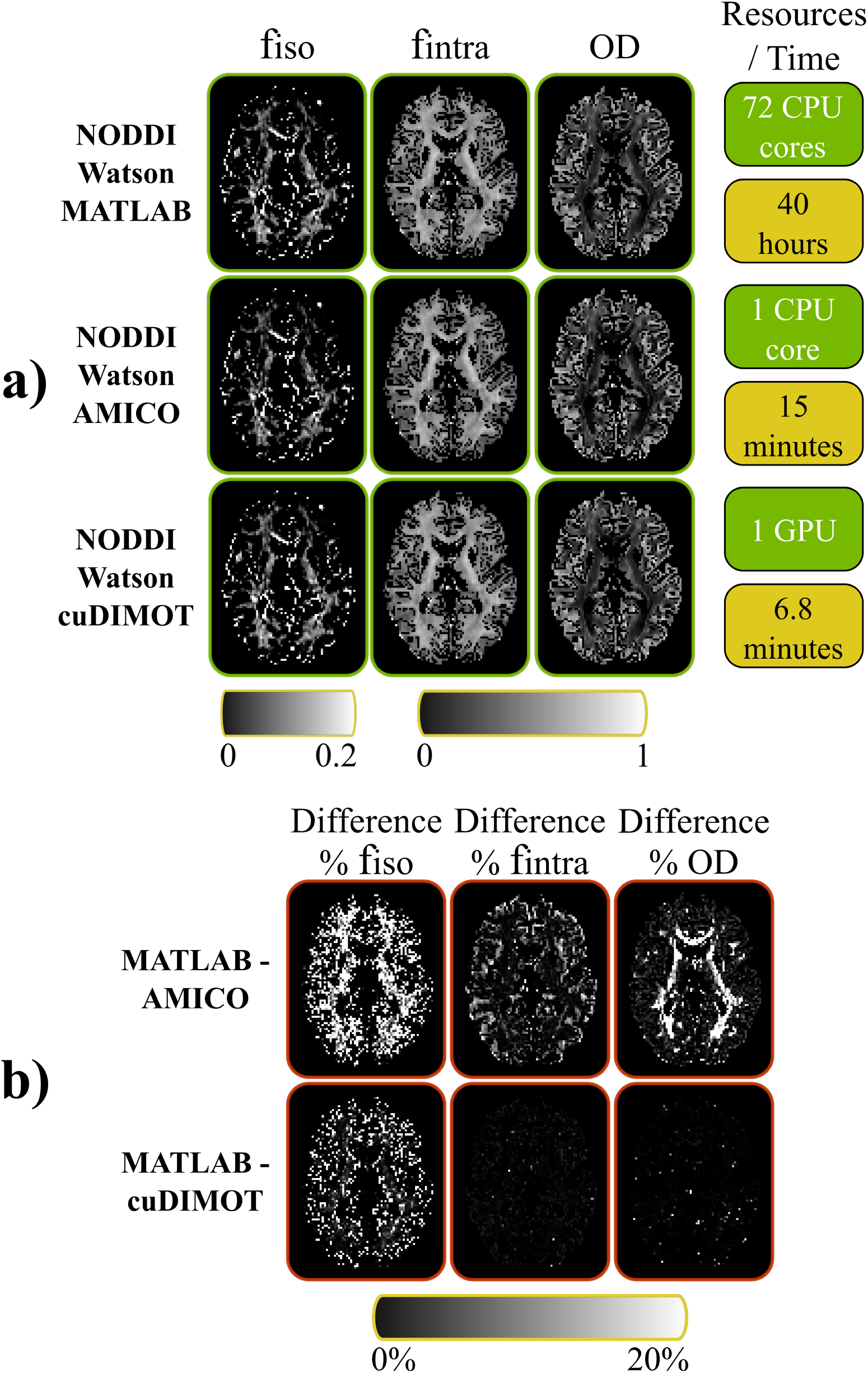
Comparison of three different tools fitting the NODDI-Watson model. (a) The results from each tool are presented in different rows. The first 3 columns show the map of the estimates for the parameters fiso, fintra and the index OD. The 4th column shows the employed computational resources and the execution times. (b) Differences, in percentage, of the estimated values between the Matlab toolbox and the other approaches.

To compare the estimates, we treat the NODDI toolbox results as ground truth, and we calculate the percentage absolute difference with the estimates obtained from the other two approaches. Figure 8b shows higher differences with AMICO than with cuDIMOT for some of the estimated parameters. The differences between the Matlab implementation and cuDIMOT are insignificant, except for the parameter f_iso_ in certain parts of the white matter. However, the values of f_iso_ are very low in the white matter, and the absolute difference between the Matlab toolbox and cuDIMOT are very small (~ 0.003) (See Figure 5 in the supplementary material). The absolute differences between the Matlab toolbox and AMICO are also small, but more significant (~ 0.03).

To further explore these findings, Figure 9 shows scatterplots of the estimated values throughout the brain. The correlations between the Matlab implementation and AMICO, and between the Matlab toolbox and cuDIMOT are presented. cuDIMOT results are highly correlated to the results from the Matlab tool. The discretisation approach used in AMICO is evident in these scatterplots, particularly for the f_intra_ parameter, where discretisation effects in the estimates are evident. For cuDIMOT, correlations with ground truth is higher. We only find some differences in a few grey matter voxels where OD takes relatively high values (OD > 0.85, i.e. very high fibre orientation dispersion). We compared the distribution of the estimated values for the parameter *κ*, from which OD is derived (See Figure 6a in the supplementary material), and we found that for low values of *κ* (*κ* <0.2), the Matlab toolbox seems to get trapped in a local minimum (*κ* = 0.5), whereas in cuDIMOT the value of the parameter is pushed towards the lower bound (κ = 0.0). Moreover, we found that this happens in a very low proportion of voxels located at the interface between grey matter and CSF (See Figure 6b in the supplementary material). We believe these differences are due to the numerical approximations used. In the cuDIMOT implementation we approximate the Dawson’s integral as in (Press et al. 1987). Likely, the Matlab toolbox is using a different approximation. Overall, these results indicate that cuDIMOT achieves very similar performance to the NODDI Matlab toolbox, and compared with AMICO, cuDIMOT is faster and obtains more accurate results.

**Figure 9.**
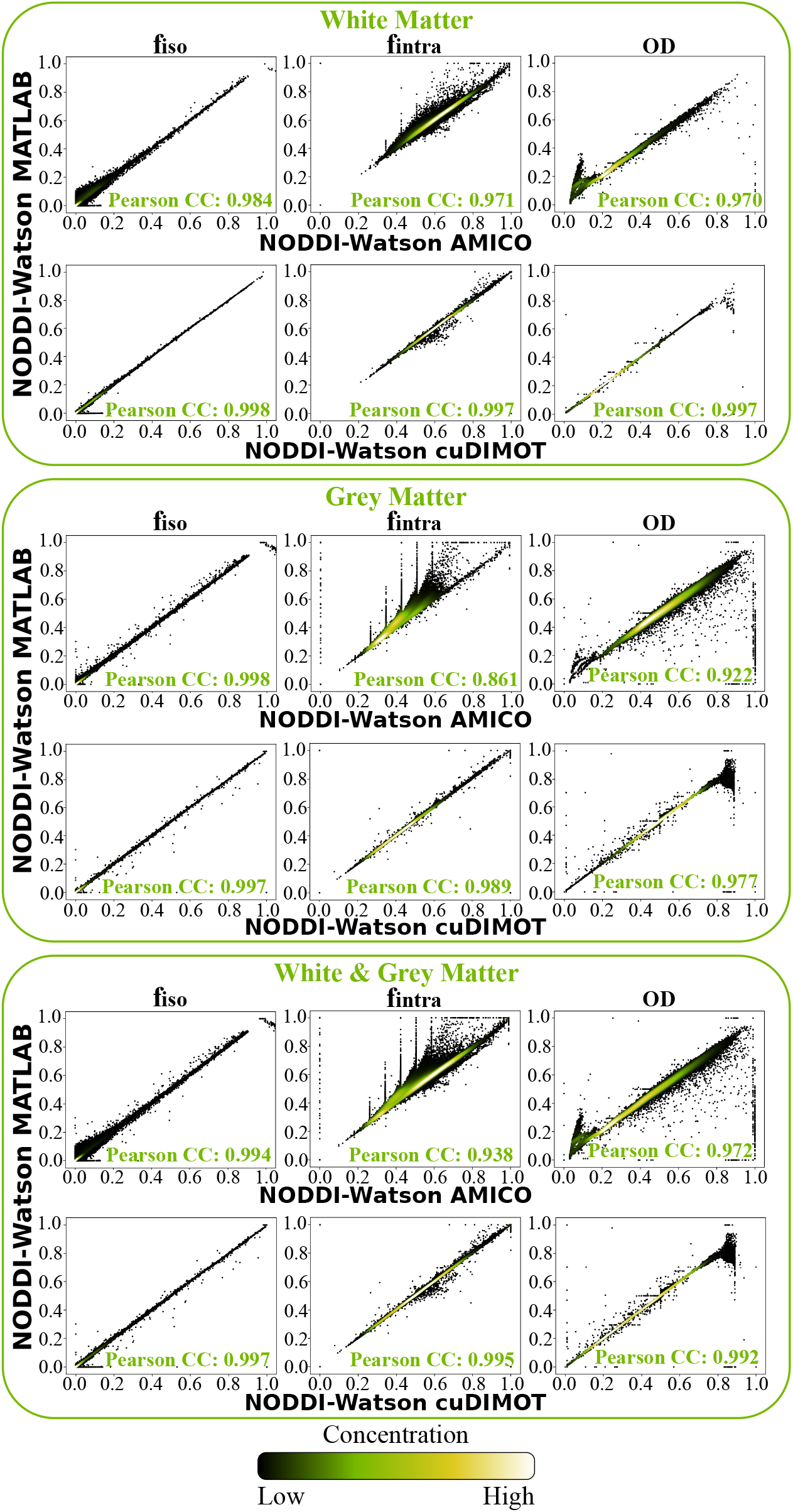
Correlations between the results from a Matlab tool and AMICO/cuDIMOT fitting the NODDI-Watson model in the white matter, grey matter and the combination of white & grey matter.

We also performed similar comparisons for the NODDI-Bingham model (Microstructure Imaging Group - University College London n.d.). Using a single GPU, cuDIMOT was found to be 7 times faster than the Matlab implementation running on a cluster with 72 cores^‡^ (Figure 10). We obtain very similar results from both tools; however, the percentage absolute differences are on average higher compared to the NODDI Watson model (bottom row). To gain further insight, figure 11 shows scatterplots of the parameter values estimated using both methods throughout the brain. In all the parameters, the correlation coefficient was higher than 0.984 in the white matter, and 0.977 in the grey matter. Notably, we found some voxels where one of the toolbox returns a very low DA (near zero) but not the other. We found that these voxels represent a very low proportion of the whole dataset, 0.2%, and they are at the interface between white matter and CSF. We believe that the source of these differences come from:

- Using a different approximation of the hypergeometric function. In cuDIMOT we use a Saddlepoint approximation (Kume & Wood 2005) and in the Matlab toolbox the function is approximated as in (Koev & Edelman 2006).
- Different non-linear optimisation method. We use Levenberg-Marquardt whereas the Matlab toolbox uses the active set algorithm included in the fmincon function (Gill et al. 1984).

**Figure 10.**
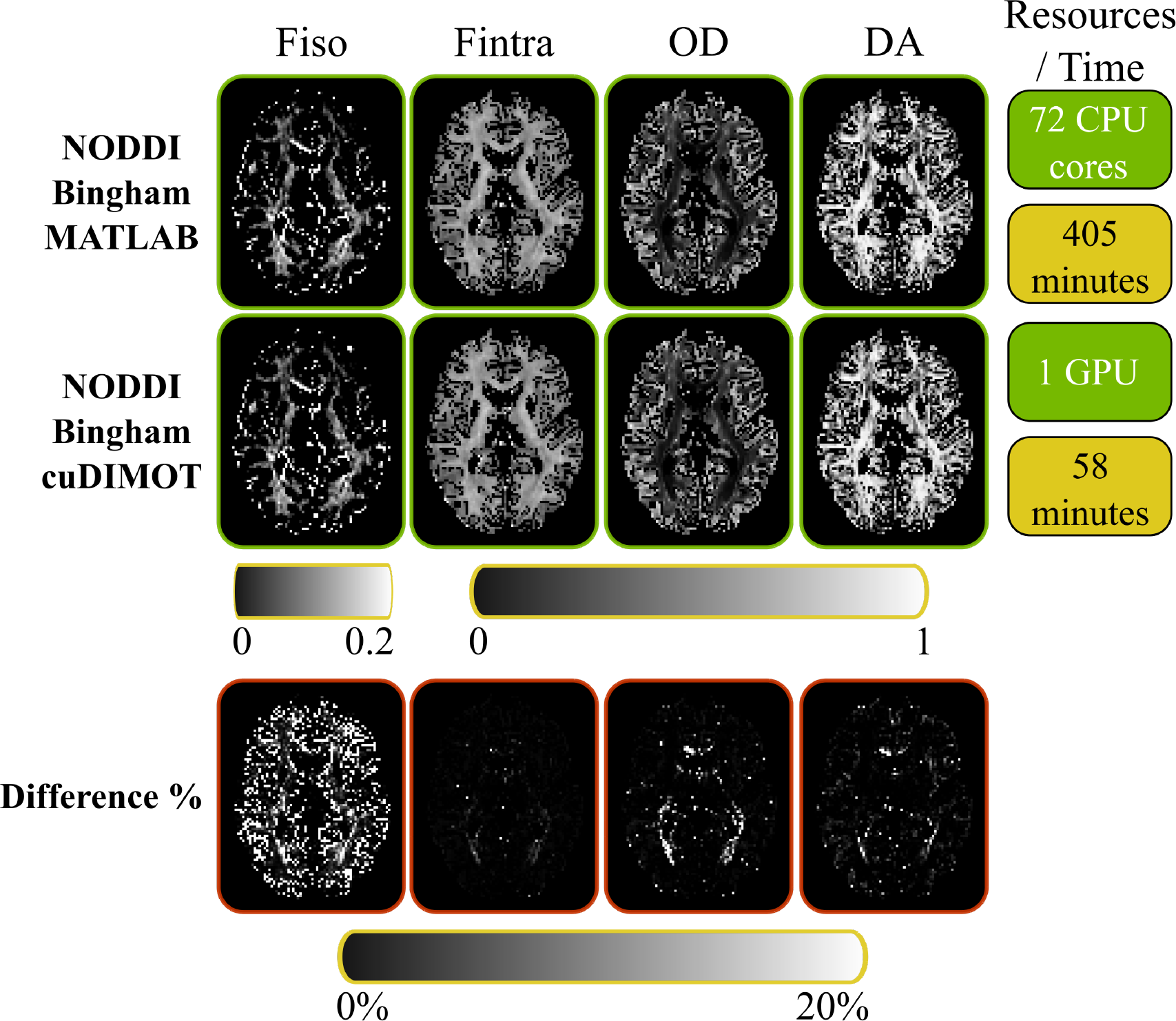
Comparison of a Matlab tool and cuDIMOT fitting the NODDI-Bingham model. The first 2 rows show the map of the estimates for the parameters fiso, fintra, the indices OD and DA, the used computational resources and execution times for each tool. The bottom row shows the differences, in percentage, of the estimated parameters between the Matlab tool and cuDIMOT.

**Figure 11.**
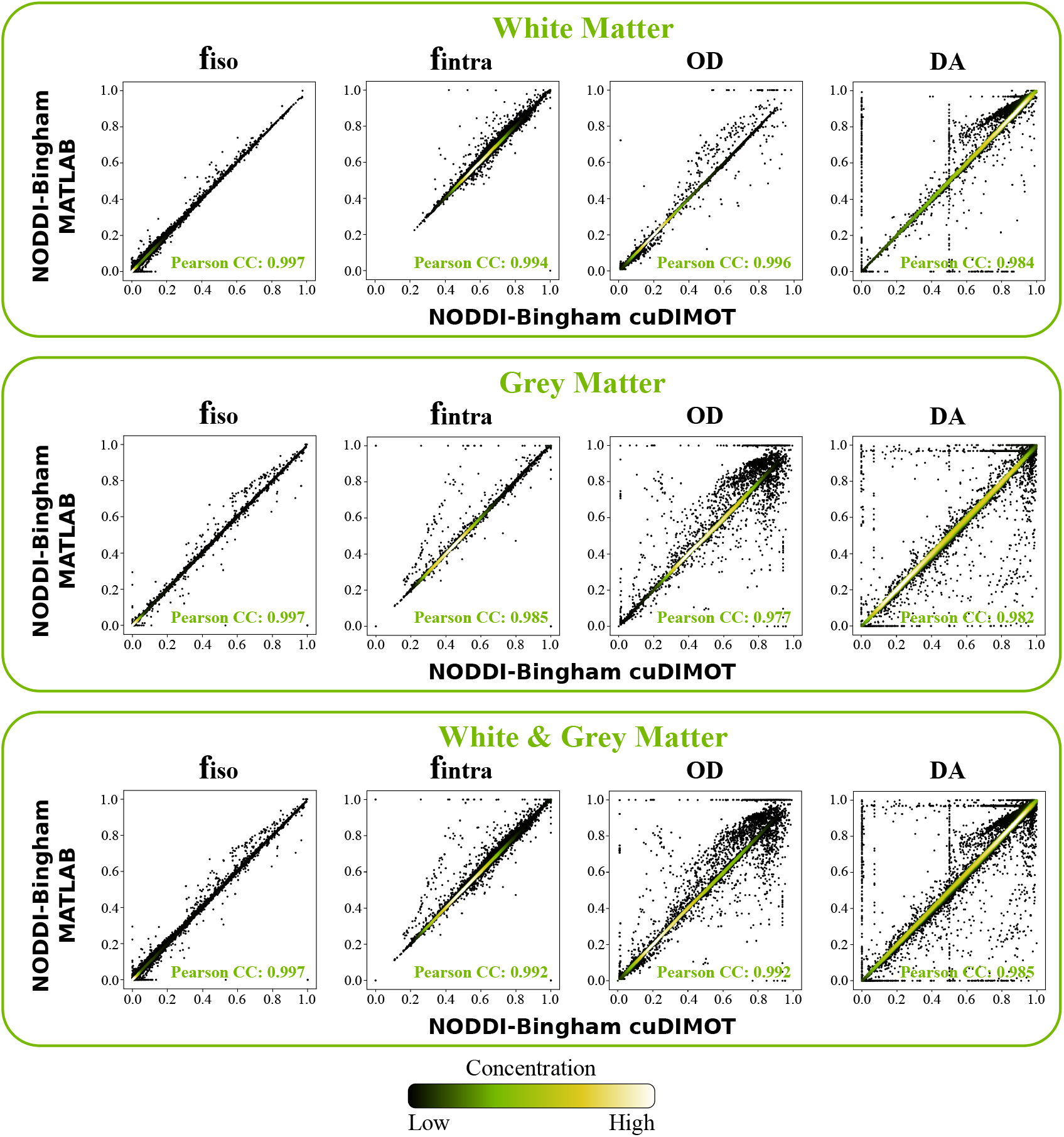
Correlations between the results from a Matlab tool and cuDIMOT fitting the NODDI-Bingham model in the white matter, grey matter and the combination of white & grey matter.

We also found a few voxels where the DA parameter is estimated with values around 0.5 in cuDIMOT, whereas in the Matlab toolbox the values are different. This seems to be related to the initial Grid-Search routine and the values that define the grid for the second concentration parameter *κ*_2_. Both Matlab toolbox and cuDIMOT reparametrize this parameter as *β* = *κ*_1_ – *κ*_2_. However, in cuDIMOT we include in the grid a set of values (from 0 to 16), whereas Matlab toolbox use a single constant value defined by the coefficient between the second and third eigenvalues of the diffusion tensor. Nevertheless, overall we obtain very high correlations between both toolboxes.

To assess speed-ups achieved by cuDIMOT, we implemented several diffusion dMRI models. On average, our toolbox achieves accelerations of two orders of magnitude compared to commonly used tools for fitting dMRI models running on a single CPU core. We report in Table 3 the speedups obtained by cuDIMOT fitting several models and using a single NVIDIA K80 device, compared to the commonly used tools running on 72 CPU cores. A Biobank dataset was used for this experiment.

**Table 3.** Speedups obtained by cuDIMOT, fitting several dMRI models to a dataset from the UK Biobank on a single K80 NVIDIA GPU, compared with the commonly used tools that implement these models and executed on a computing cluster and using 72 CPU cores (and a single CPU thread per core).

|  | Ball<br>& 1-stick | Ball<br>& 2-sticks | Ball<br>& 1-stick +<br>Gamma | Ball<br>& 2-sticks +<br>Gamma | NODDI<br>Watson | NODDI<br>Bingham |
| --- | --- | --- | --- | --- | --- | --- |
| Common Tools<br>72 CPU cores | 720s | 1380s | 1260s | 2520s | 2400m | 405m |
| cuDIMOT<br>single NVIDIA K80<br>device | 187s | 423s | 324s | 679s | 6.8m | 58m |
| Speedup | 3.85x | 3.26x | 3.88x | 3.7x | 352x | 6.98x |

Compared to a C++ implementation run on the computing cluster, (Jenkinson et al. 2012) cuDIMOT is on average 3.66 times faster. We considered the following models:

- Ball & 1 stick (Behrens et al. 2003; Behrens et al. 2007)
- Ball & 2 sticks
- Ball & 1 stick (with gamma-distribution for the diffusivity (Jbabdi et al. 2012))
- Ball & 2 sticks (with gamma-distribution for the diffusivity)
- NODDI-Watson
- NODDI-Bingham

To illustrate the flexibility of cuDIMOT in defining new models and the benefit from accelerations that allow extensive model comparison, even with stochastic optimisation, we used cuDIMOT to test whether crossing or dispersing models are more supported by the data. We performed a comparison of six diffusion MRI models and used the BIC index for comparing the performance. The models included in this test were:

- Ball & 1 stick (with gamma-distribution for the diffusivity (Jbabdi et al. 2012))
- Ball & 2 sticks (with gamma-distribution for the diffusivity)
- NODDI-Watson
- Ball & racket (Sotiropoulos et al. 2012)
- NODDI-Bingham
- NODDI-2-Binghams: we implement an extension of the NODDI-Bingham model (Tariq et al. 2016) for including two fibre orientations with the model signal given by:

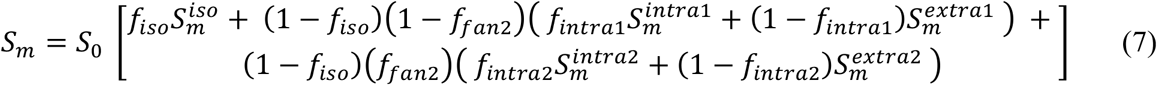

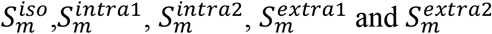 are defined as in the NODDI-Bingham model.

The model has a total of 14 free parameters:

- Compartments fraction: *f_iso_*, *f_fan_*_2_, *f_intra_*_1_, *f_intra_*_2_
- First fibre distribution: *κ*_1_1_, *κ*_1_2_, *θ*_1_, ϕ_1_, *ψ*_1_
- Second fibre distribution: *κ*_2_1_, *κ*_2_2_, *θ*_2_, ϕ_2_, *ψ*_2_

In all cases we ran an initialisation routine (Grid-Search or the output of the fitting process of another model), we run Levenberg-Marquardt and MCMC. cuDIMOT calculates the BIC from the mean of the parameter estimates. We first classify the six models into two groups, one group with the models that do not characterise the dispersion of fibre orientations, which include the ball & stick(s) models, and another group with the models that characterise the dispersion. The second row in Figure 12 shows a colour-coded map indicating in what voxels each group gets a better BIC (lower), i.e. its complexity is better supported by the data. Using dispersion models the diffusion signal is better explained, and the obtained BIC is lower in the majority of brain regions. The last row of Figure 12 compares the four considered dispersion models. The dominant model that gets a lower BIC is NODDI-Bingham (55% of the voxels) followed by NODDI-Watson (24% of the voxels), consistent with the results presented by (Ghosh et al. 2016). Interestingly, 5% of the voxels particularly in the centrum semiovale support the existence of dispersing populations crossing each other.

**Figure 12.**
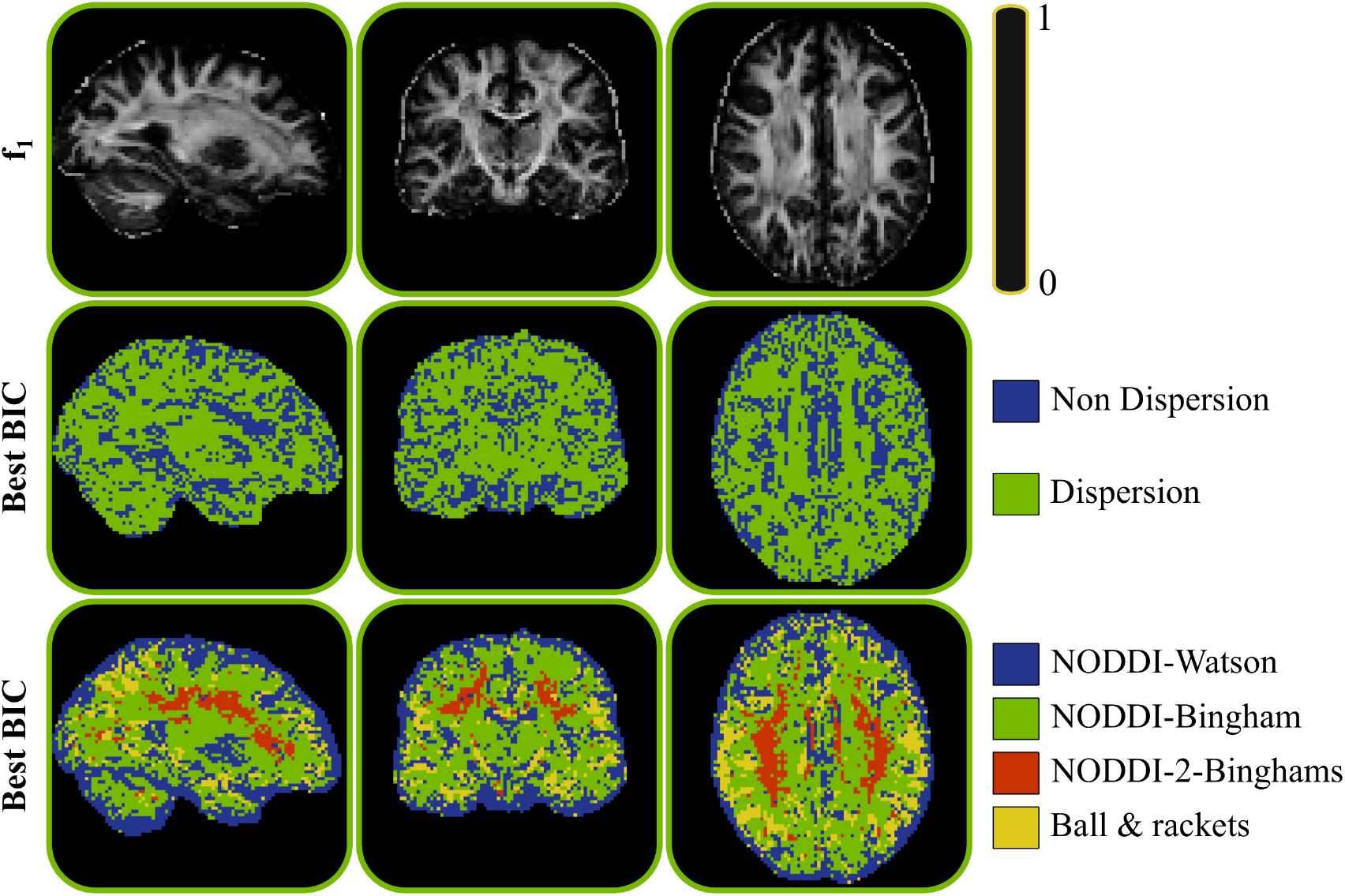
Model performance comparison. The first row shows a map for reference with the estimated fraction of the principal fibre in the ball & 2 sticks model. The second and third rows show color-coded maps indicating in what locations a model or a group of models get the best BIC.

### Tractography with GPUs

In order to validate the GPU-accelerated probabilistic tractography framework we performed various tests and compared the results with the results obtained using a CPU-based tractography application (Behrens et al. 2007; Smith et al. 2004), as implemented in FSL, for both white matter tract reconstruction and connectome generation. Given the stochastic nature of probabilistic tractography methods, we expect some variability in the results of both frameworks, but given the high number of samples that we have used in the tests, we expect the results to have converged and the variability to be small. Nevertheless, we run every experiment 10 times and we compare the differences between CPU and GPU results with the run-rerun variability.

Figure 13a shows some quantitative comparisons of reconstructed tracts using both implementations. We reconstructed 27 major white matter tracts as in (de Groot et al. 2013) (12 bilateral and 3 commissural ones) using standard space tracking protocols and constraints (see Table 4 for a list of reconstructed tracts). In a different test, we generated dense connectomes using the HCP grayordinates (91K seed points (Glasser et al. 2013)). To quantify these comparisons, we present the run-rerun the variability of each framework independently and the distribution of correlation coefficients between the CPU-based and the GPU-based frameworks. In the reconstruction of the 27 tracts the correlation was calculated voxel-wise. In the generation of the dense connectome, the correlation was calculated from all the elements of the connectivity matrix. The individual run-rerun correlation coefficients are higher than 0.999 in all the cases, for both the CPU and the GPU frameworks. Importantly, the correlation coefficients between CPU and GPU are higher than 0.998, illustrating that the two implementations provide the same solution. Even if these correlations are slightly lower than between the individual run-rerun results (CPU vs. CPU and GPU vs. GPU), this is expected, as some mathematical operations have different implementations (e.g. rounding modes) and different precision in a GPU compared with a CPU (Whitehead & Fit-florea 2011). Figure 13b shows a qualitative comparison of the reconstruction of six exemplar tracts using both frameworks.

**Table 4:**
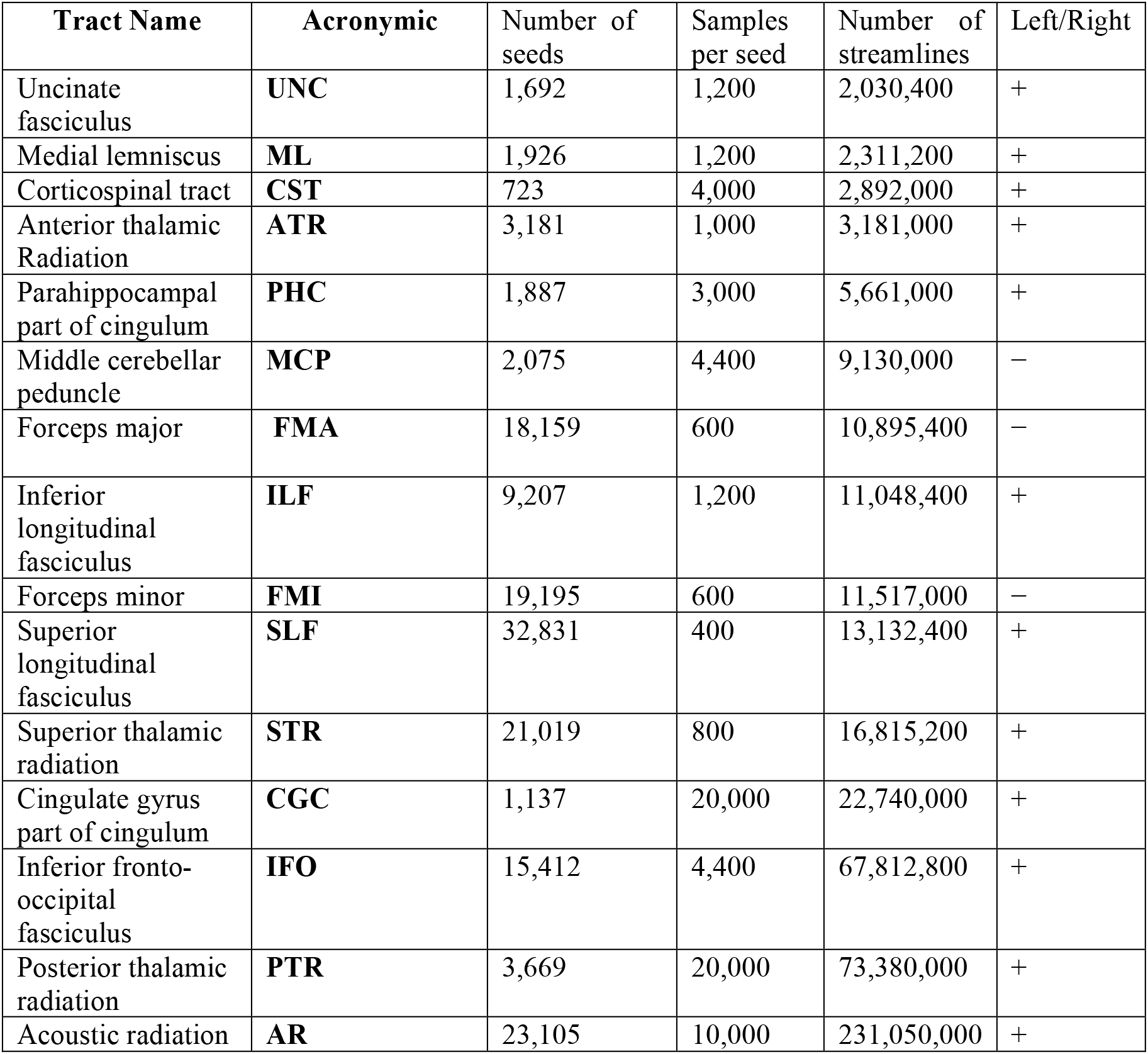
List of reconstructed tracts sorted by number of propagated streamlines. Different number of seed points and samples are used for reconstructing the tracts. Some tracts have a bilateral homologue (+) and some others no (−).

| Tract Name | Acronymic | Number of seeds | Samples per seed | Number of streamlines | Left/Right |
| --- | --- | --- | --- | --- | --- |
| Uncinate fasciculus | **UNC** | 1,692 | 1,200 | 2,030,400 | + |
| Medial lemniscus | **ML** | 1,926 | 1,200 | 2,311,200 | + |
| Corticospinal tract | **CST** | 723 | 4,000 | 2,892,000 | + |
| Anterior thalamic Radiation | **ATR** | 3,181 | 1,000 | 3,181,000 | + |
| Parahippocampal part of cingulum | **PHC** | 1,887 | 3,000 | 5,661,000 | + |
| Middle cerebellar peduncle | **MCP** | 2,075 | 4,400 | 9,130,000 | – |
| Forceps major | **FMA** | 18,159 | 600 | 10,895,400 | – |
| Inferior longitudinal fasciculus | **ILF** | 9,207 | 1,200 | 11,048,400 | + |
| Forceps minor | **FMI** | 19,195 | 600 | 11,517,000 | – |
| Superior longitudinal fasciculus | **SLF** | 32,831 | 400 | 13,132,400 | + |
| Superior thalamic radiation | **STR** | 21,019 | 800 | 16,815,200 | + |
| Cingulate gyrus part of cingulum | **CGC** | 1,137 | 20,000 | 22,740,000 | + |
| Inferior fronto-occipital fasciculus | **IFO** | 15,412 | 4,400 | 67,812,800 | + |
| Posterior thalamic radiation | **PTR** | 3,669 | 20,000 | 73,380,000 | + |
| Acoustic radiation | **AR** | 23,105 | 10,000 | 231,050,000 | + |

**Figure 13.**
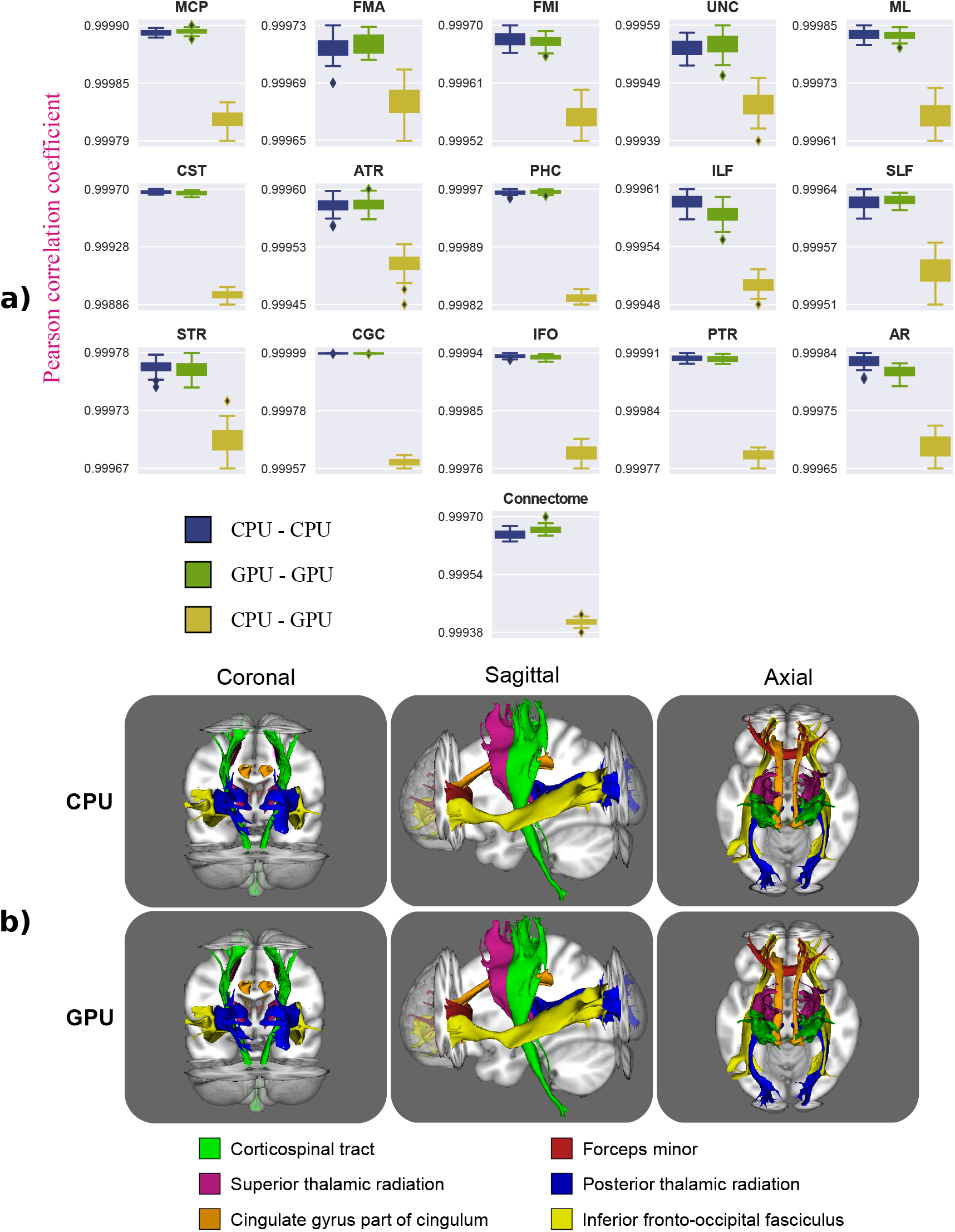
(a) Run-rerun variability of CPU-based and GPU-based probabilistic tractography frameworks and distribution of the correlation coefficients between both frameworks. Results are showed in the reconstruction of 12 bilateral tracts, 3 commissural tracts and in the generation of a dense connectome. Each experiment was run 10 times. The 45 combinations of correlations between re-runs were considered and 45 out of the 100 combinations of CPU vs. GPU correlation coefficients were chosen randomly. (b) Coronal, sagittal and axial views comparing CPU-based and GPU-based frameworks performing probabilistic tractography and reconstructing some major white matter tracts. Each colour represents a different white matter tract. These paths are binarised versions of the path distributions after being thresholded at 0.5%.

We evaluated the optimal number of CUDA streams (and OpenMP threads) for each test (see Figure 14). The most efficient configurations were 8 CUDA streams for generating dense connectomes, obtaining a total gain of 1.35x respect using 1 CUDA stream, and 4 CUDA streams for reconstructing tracts, obtaining gains of only 1.12x. Figure 15 reports computation times reconstructing the 12 bilateral tracts and the 3 commissural tracts individually. A single CPU core was used for running the CPU-based framework, and a single GPU and four CUDA streams for processing the tracts with the GPU-accelerated framework. On average, a speedup of 238x was achieved, in the range of 80x to 357x. In all cases, except for the reconstruction of the Acoustic Radiation (AR), the GPU-based application achieves accelerations of more than two orders of magnitude. In general, if the reconstruction of a tract involves several anatomical constraints that makes the algorithm to stop or discard streamlines at early steps, including tight termination and exclusion masks, the GPU-based framework performs worse, as these masks are not checked until the propagation of the streamlines has completely finished (see Figure 7a). The reconstruction of the Acoustic Radiation uses a very tight exclusion mask and thus the achieved performance is lower compared with the reconstruction of other tracts.

**Figure 14.**
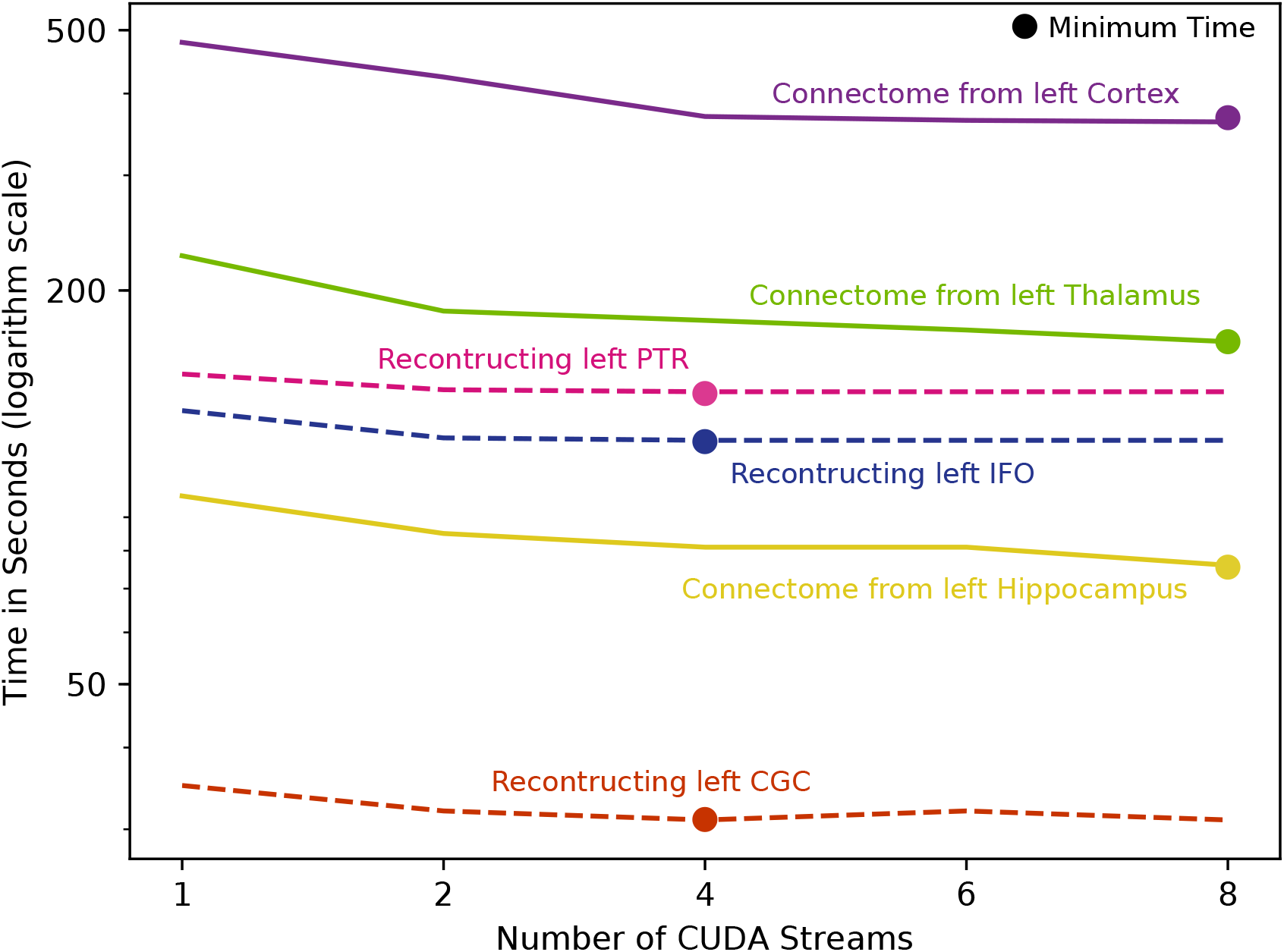
Execution times of the GPU tractography framework using different numbers of CUDA streams. Results are shown when the application reconstructs three different tracts in the left hemisphere (Inferior fronto-occipital fasciculus, Posterior thalamic radiation and Cingulate gyrus part of cingulum), and when the application generates a connectivity matrix using three different regions as seeds (left Cortex, left Thalamus and left Hippocampus).

**Figure 15.**
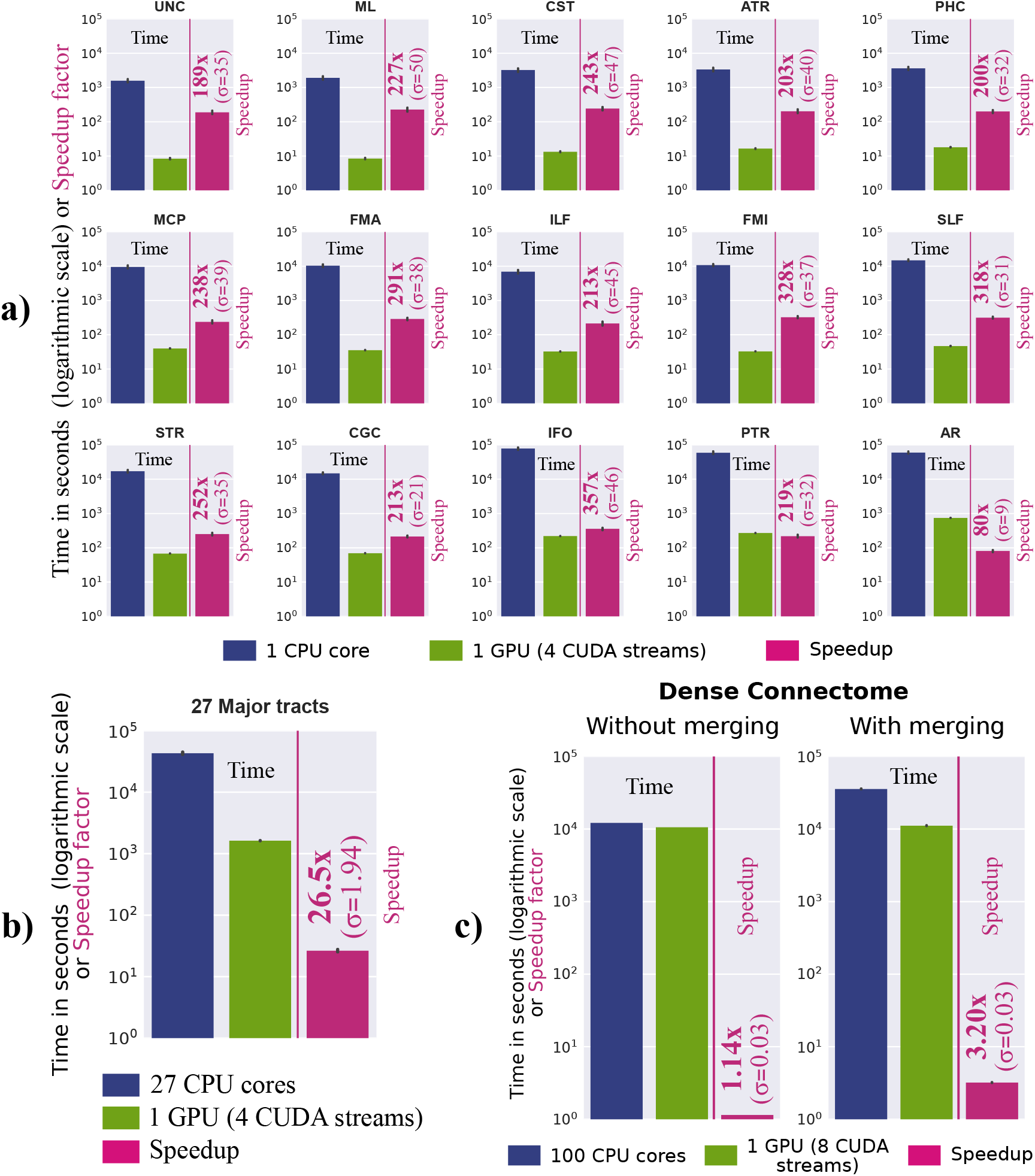
(a) Execution times (in logarithmic scale) and speedup (standard deviation σ is also shown) in the reconstruction of 12 bilateral tracts and 3 commissural tracts comparing a GPU-based with a CPU-based probabilistic tractography framework. Execution times (in logarithmic scale) and speedup (and its standard deviation σ) reconstructing 27 tracts are presented in (b) and generating a dense connectome in (c).

Figure 15b reports the total execution time reconstructing all the tracts. When the CPU-based tool is used, the reconstruction of several tracts can be parallelised. Tracts are completely independent and thus their reconstruction can be processed by different threads. A total of 27 CPU cores were used in this case, using different CPU threads for reconstructing different tracts. A single GPU and four CUDA streams were used again for processing the tracts with the GPU-accelerated framework, processing sequentially the different tracts. A speedup of 26.5x was achieved using the GPU-accelerated solution.

We use the CPU-based and the GPU-based frameworks for generating a dense connectome. 91,282 seed points and 10,000 samples per seed point were employed, having a total of 912.82 million streamlines. For generating the connectome with the CPU-based application we used 100 CPU cores, each one propagating 100 streamlines from each seed point. This process took on average 3.38 hours. At the end of the process, the different generated connectomes (on different CPU cores) need to be added. This merging process took on average 6.5 hours (due to the size of the final connectivity matrices).

We used 1 single GPU and 8 CUDA streams for generating the connectome with the GPU-based application. The process took on average 2.95 hours. Figure 15c reports these execution times and the speedup achieved by the GPU-based framework, with and without considering the merging process required by the CPU multi-core application. Without considering the merging process both applications reported similar execution times. Considering the merging process, the GPU application was more than three times faster than the CPU multi-core application. In both cases, 1 GPU offers at least two orders of magnitude speed-up compared with a single CPU core.

Apart from the computational benefits, we have added new functionality to the GPU tractography toolbox. A novel feature is the possibility of using surfaces for imposing more accurate anatomical constraints. For instance, we can use a pial surface as termination mask for avoiding the propagation of streamlines outside this surface, and avoiding non-plausible connections along the CSF in the subarachnoid space. As shown in the results of Figure 16a, surfaces allow us to describe more accurately the cortical folding patterns and allow more accurate constraints to be imposed.

**Figure 16.**
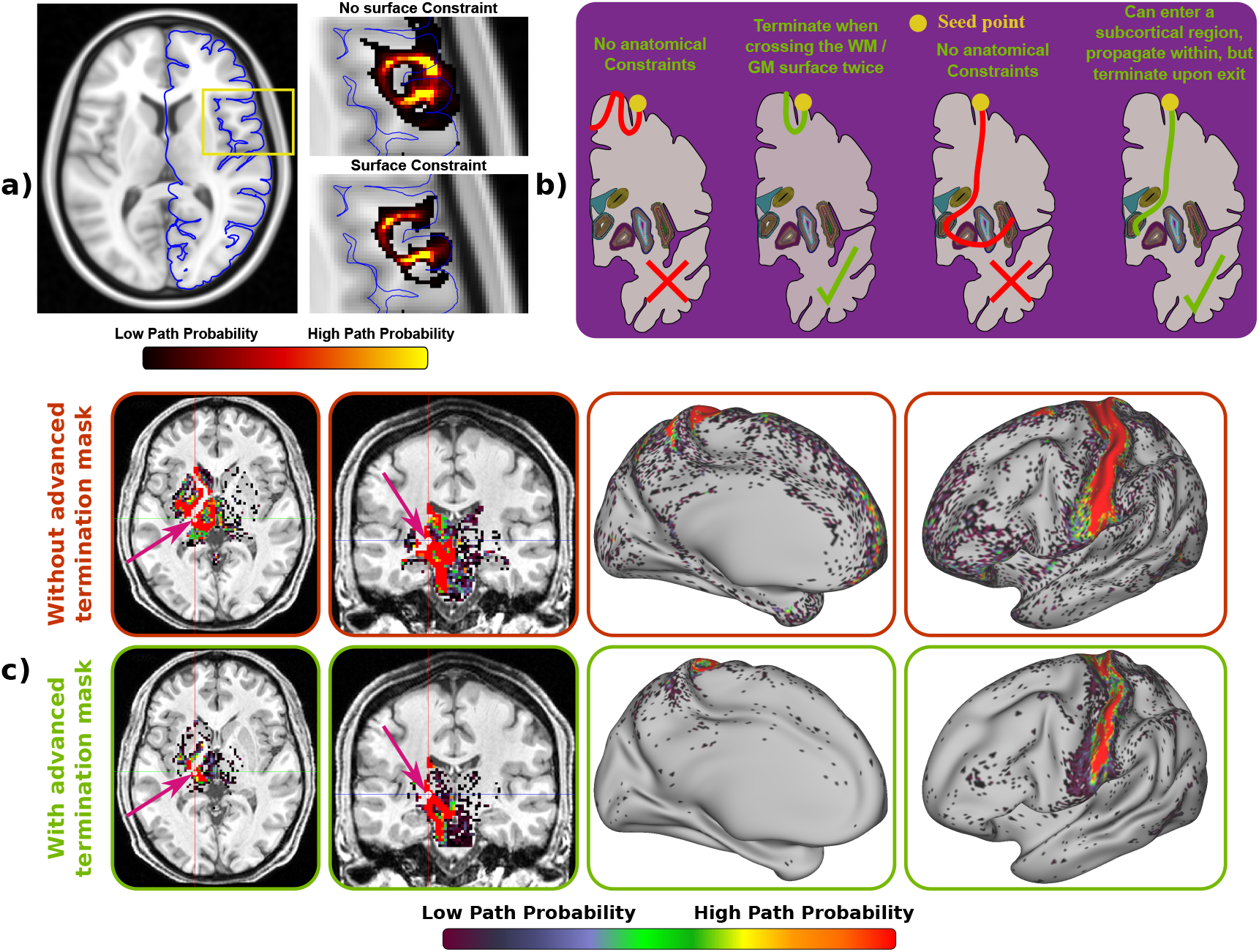
(a) Example of the use of surfaces for imposing anatomical constraints. Probabilistic tractography is performed using as seed and target points the right inferior frontal gyrus. Without using a surface constraint, wrong paths that jump between neighbouring gyri can be generated. (b) Advanced termination masks. The tractography framework adds the possibility of stopping the streamlines when they cross a surface twice and/or streamlines can be propagated inside a subcortical region but the framework stops them upon exit. (c) Connectivity from a voxel inside the left Thalamus (fuchsia arrow) using and not using advanced terminations masks. The first two columns show the connectivity with other subcortical structures. The last two columns show the connectivity with all the vertices on the left hemisphere cortex.

A more sophisticated termination mask mechanism has also been added to the GPU framework. Commonly, termination masks force the algorithm to stop the propagation of the streamlines the first time they hit the mask, but sometimes it is reasonable to allow the propagation until certain conditions are met (see Figure 16b). For instance, to increase the chances of getting “direct” connections, it is desired that a streamline crosses the WM/GM boundary no more than twice when reconstructing corticocortical networks, once at the starting point and once at the end point. However, it does not seem plausible to have pathways running in and out of the WM/GM boundary or in parallel along the cortex connecting several regions. Thus, a special type of termination masks can be used for stopping the streamlines when they cross a surface twice. Similarly, to encourage direct cortico-subcortical connections, it is undesirable that a streamline visits several subcortical regions, but ideally we would like a streamline to be able to propagate within a subcortical region. As in (Smith et al. 2012), our framework can use the special termination masks for stopping the streamlines upon exiting these regions, while allowing propagation within them. Figure 16c shows the effect of imposing these anatomical constraints when generating a dense connectome. The seed mask is defined with a WM/GM boundary surface and termination volumes include several subcortical structures (accumbens, amygdala, caudate, cerebellum, hippocampus, pallidus, putamen and thalamus). The connectivity pattern from the sensori-motor part of the thalamus without and with advanced termination masks is illustrated. In the former case, the streamlines can cross the cortex or subcortical structures several times and continue propagating, generating a number of false positives (for instance see hotspots along the frontal medial surface). In the latter case, this situation is avoided, and a more realistic connectivity map is obtained, connecting the sensorimotor part of the thalamus to sensorimotor cortical regions.

## DISCUSSION

We have presented GPU-based parallel computational frameworks for accelerating the analysis of diffusion MRI, spanning from voxel-wise biophysical model fitting to tractography and whole-brain connectome generation. Despite the difference in the inherent parallelisability of these applications, GPUs can offer considerable benefits when challenges are carefully considered. Speed-ups of over two orders of magnitude change the perspective of what is computationally feasible. The GPU toolboxes will be publically released as part of FMRIB’s Software Library (FSL).

The accelerations achieved by the designs proposed here can be tremendously beneficial. Big databases arise more and more often from large consortiums and cornerstone projects worldwide. Hundreds or even thousands of datasets need to be processed. The throughput of the parallel designs using a single or a multi-GPU system is higher than a CPU multi-core system. Very large recent studies such as the Human Connectome Project (HCP) (Van Essen & Ugurbil 2012; Van Essen et al. 2012; Sotiropoulos et al. 2013), (data from 1,200 adults), the Developing Human Connectome Project (dHCP) (data from 1,000 babies) and UK Biobank (Miller et al. 2016; Alfaro-Almagro et al. n.d.) (data from 100,000 adults) are using our parallel designs for processing these datasets on GPU clusters (Hernández et al. 2013). For instance, a 10-GPU cluster has been built for processing the most computationally expensive tasks of the UK Biobank pipeline. The cluster allows fitting the ball & sticks model to 415 datasets per day. Running the same tasks with a cluster of 100 CPU cores, only 25 datasets could have been processed per day. Moreover, to obtain a similar throughput as the 10-GPU cluster, more than 1,600 CPU cores would have been necessary.

Apart from increasing feasibility in big imaging data exploration, the designs presented here can assist in model exploration and development, as estimation and testing is an inherent part of model design. Moreover, the response time for analysing a single dataset is dramatically reduced from several hours/days to few minutes using a single GPU, and close to real-time processing could make these methods more appealing for clinical practice.

### GPU-based biophysical modelling

We have presented a generic modelling framework, cuDIMOT, that provides a model-independent frontend, and automatically generates a GPU executable file that includes several fitting routines for user-defined voxel-wise MRI models. We have used cuDIMOT to explore diffusion models that characterise fibre orientation dispersion, and we have shown that it can be very useful for exploring, designing and implementing new dMRI protocols and models. It is easy to use and generates very efficient GPU-accelerated solutions.

A toolbox with the same purpose as cuDIMOT, has been recently presented (Harms et al. 2017). However, it does not include the option for performing stochastic optimisation or the ability to add priors on the model parameters. Furthermore, its design only includes the first level of parallelisation where the voxel fitting process of different voxels are distributed among threads, leading to lower speed-ups (accelerations from 11X to 38X comparing a GPU with a single CPU core were reported in (Harms et al. 2014)). It is implemented in a more generic (non-GPU specific) programming model (OpenCL (Stone et al. 2010) rather than CUDA), allowing the parallelisation on both multi-core CPUs and GPUs, but achieving lower performance on NVIDIA GPUs.

Our toolbox has been designed for processing datasets with any number of voxels and measurements. If the memory required for storing the data of all the voxels exceeds the device memory, the framework divides the data into several subsets (according to the amount of device memory available), and these subsets are processed one after the other. However, before being processed, each subset needs to be copied into device memory. This can create a performance penalty if the device memory capacity is small (1 or 2 GB) because a large number of CPU to GPU transfers is required, and these transfers are expensive (using normally the PCIe interconnection bus). This is however not a problem in the new NVIDIA architectures, where global memory space is larger (up to 24 GB in Pascal architecture (NVIDIA 2016)) and a new CPU-GPU interconnection bus is incorporated (NVLINK (NVIDIA 2014b)).

There is a limitation in cuDIMOT on the number of parameters of a model. In the Levenberg-Marquardt routine the number of model parameters is limited to 31. The cause of this limitation is in the implementation of the LU solver (see Figure 4), where each thread of a warp processes a column of the matrix for solving the system. For a model with P parameters, P + 1 threads are required, and a warp has 32 threads. In the MCMC routine there is also a limitation on the number of model parameters. The framework stores in Shared memory the parameters and some associated information (priors, number of proposals accepted/rejected, and standard deviation of the proposals distribution). Thus, a model is limited to a maximum number of around 300 parameters (the exact number depends on the size of Shared memory of the specific GPU and on the precision used to store the parameters, single or double).

A large number of dMRI modelling approaches have been proposed in the literature, but it seems that no single approach can explain all complex microstructure patterns (Ghosh et al. 2016; Ferizi et al. 2015). Thus, applications that consider several models for selecting the best one in each voxel seems to be a potential solution. Given the computational cost of fitting these models, parallel solutions like cuDIMOT will be essential for performing this type of analysis. We believe that cuDIMOT is going to be very useful in the development and improvement of new diffusion MRI models, which may explain the complexity of the diffusion process, extract useful biophysical parameters and contribute to the development of new biomarkers.

### GPU-based tractography

We have also developed and presented a probabilistic tractography framework that achieves accelerations of more than two orders of magnitude compared to a sequential solution, and can handle situations ranging from simple white matter tracking tasks to dense connectome generation. The implementation offers the possibility of defining tractography protocols with either volumes or surfaces, and the possibility of using advanced termination masks that allow more accurate anatomical constraints. We have shown the benefits of using this extended functionality.

Tractography algorithms pose particular challenges for designing GPU solutions. Each GPU thread accesses different memory locations during its execution, and these accesses cannot be anticipated, as the propagation movements are decided on the fly. Moreover, given the stochastic nature for choosing the orientation samples, the threads may diverge even if their streamlines are initialised from the same seed point. This behaviour leads to uncoalesced memory accesses and imbalanced execution length of the threads, and consequently, to a waste of GPU resources.

We studied the execution length distribution across streamlines reconstructing the same tract. In many scenarios, many generated streamlines terminate relatively quickly (before 100 steps), as they meet a termination criterion. However, there are other streamlines that take considerably more steps. This trend has also been confirmed before in [17] and [57], and it is also supported by the underlying anatomy: the majority of white matter connections are short in length (Donahue et al. 2016). To avoid a waste of resources, we explored the approach proposed in (Xu et al. 2012; Mittmann et al. 2008), where the kernel that propagates the streamlines is stopped after a certain number of steps and the threads that are idle, i.e., the threads with terminated streamlines, are removed. When the kernel is launched again on the GPU, there will be only threads with streamlines still propagating, and thus, the device resources will be used more efficiently. This also allows other streamlines to start to be processed, as memory resources are freed after removing terminated streamlines. We tried several strategies for deciding the number of propagation steps to use before stopping the kernel. The process for removing idle threads is executed on the host and it has some extra cost that may cancel out the gains of this approach. Contrary to suggestions in (Xu et al. 2012; Mittmann et al. 2008), where gains of 4x were reported when using streamlines removal strategies, we could not find enough supporting evidence that this approach results in significant performance gains. This strategy barely reduces execution times in our framework, and in some cases, it even increases them. We believe that the extended and more complex functionality offered by our tractography framework, compared with previous designs, is the main reason for these differences. Given the complex functionality, more information needs to be stored and reloaded by the GPU threads every time that the CUDA kernel is stopped, causing a higher overload. These previous studies have reported final performance gains in the range of 40x-50x, which is considerably lower than what we found here.

Another challenging requirement of the parallel tractography application is the large amount of required memory. Given the non-predictable path and length of the streamlines, all the orientation samples and memory for storing the maximum possible number of visited coordinates need to be allocated. The GPU global memory is used for storing this data. This restricts the number of streamlines that can be propagated in parallel. However, most modern GPUs have at least 3 GB of global memory, and they still can run a considerable number of streamlines (~ 40,000) in parallel. When a more complex and demanding functionality is used, such as generating a dense connectome, this number can be reduced by 60% (as more data per streamline is needed) and GPUs with at least 5 GB should be used for achieving a good performance.

Given the speed and facility to run-rerun probabilistic tractography using the developed GPU toolbox, we performed a convergence study. We evaluated the number of samples that are needed per seed point when generating a dense connectome in order to achieve convergence. To do that, we generated a dense connectome multiple times using a different number of samples per seed point. Figure 17 shows the correlation coefficients with respect to an assumed converged dense connectome, which was generated using 100,000 samples per seed. The figure also shows the correlation coefficients between consecutive runs in terms of number of samples per seed. It seems that even with 1,000 samples and 18 minutes run, the results are almost converged. Using 10,000 samples per seed achieves convergence, while the time for generating the connectome is still reasonable, less than 3 hours using a single GPU and 8 CUDA streams.

**Figure 17.**
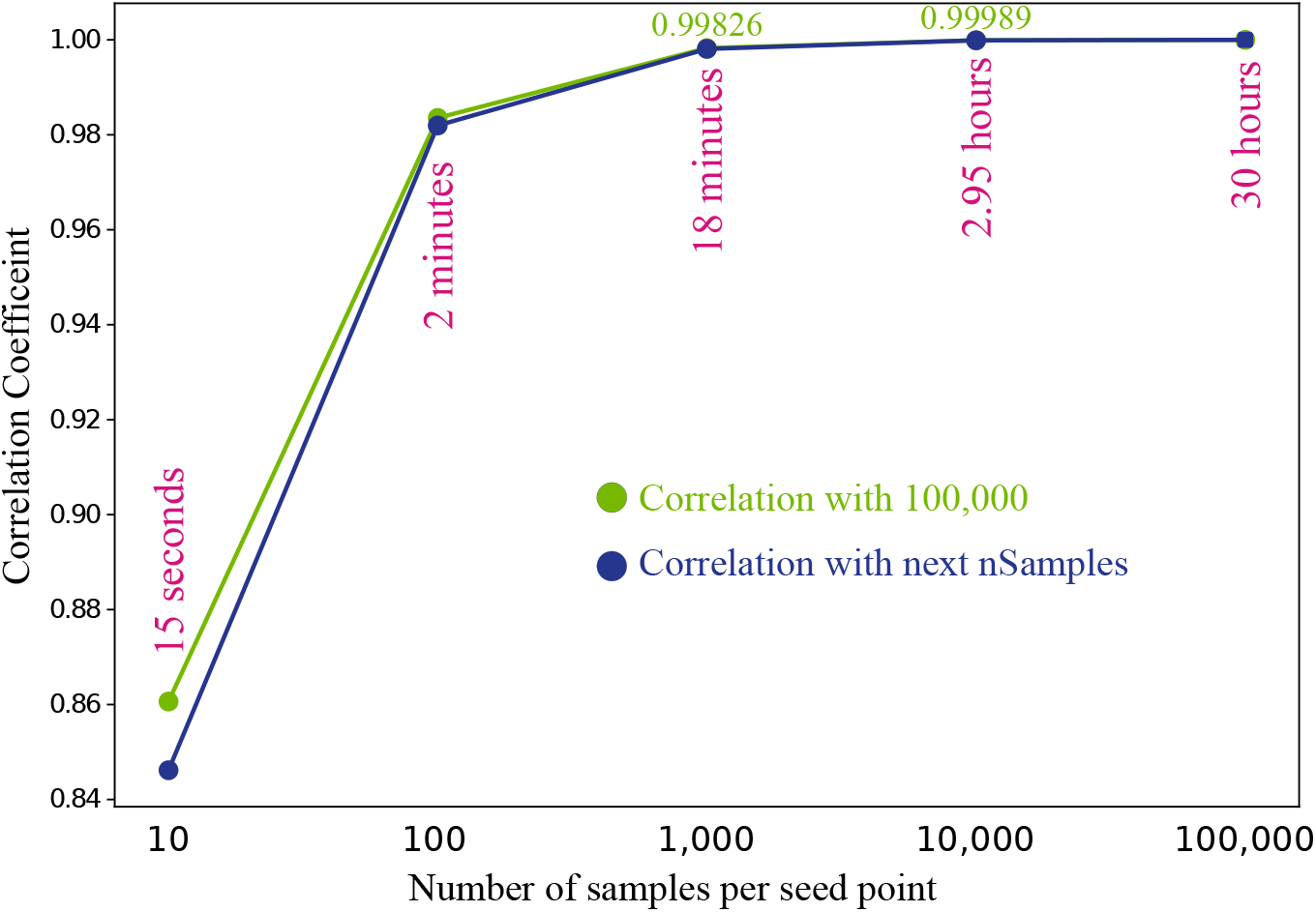
Correlation coefficient, generating a dense connectome (all greyordinates to all greyordinates) on the GPU-based framework, between re-runs, modifying the number of samples per seed point. The figure reports the correlation coefficients with respect to a dense connectome generated with 100,000 samples (green) and with respect to the connectome generated with the next number of samples in the plot (blue). The figure also shows the execution times generating these dense connectome on a single NVIDIA K80 GPU.

A final consideration is that the designs presented here have been optimised for a specific type of GPUs, NVIDIA Tesla K80. Although the frameworks works for any NVIDIA GPU (with double precision capability), the optimal number of CUDA threads per block and the distribution of voxels amongst the CUDA blocks may be different for other types of GPU architectures, such as Pascal (NVIDIA 2016). Options for autotuning and auto-compilation (Giles et al. 2013) could be integrated with the framework, but the evaluation of these is left for future work. Nevertheless, accelerations of more than two orders of magnitude have been reported by many other users of our toolboxes using different NVIDIA architectures.

## ACKNOWLEDGMENTS

We would like to acknowledge financial support from the UK Engineering and Physical Sciences Research Council (EP/L023067/1). Part of this project was awarded the NVIDIA 2016 GPU centre of excellence achievement, and the prize was used for partially funding this research. We also acknowledge the support of NVIDIA Corporation with the donation of the Titan X Pascal GPU used for aspects of development of the presented toolboxes. Data were provided by the Human Connectome Project, WU-Minn Consortium (Principal Investigators: David Van Essen and Kamil Ugurbil; 1U54MH091657) funded by the 16 NIH Institutes and Centers that support the NIH Blueprint for Neuroscience Research; and by the McDonnell Center for Systems Neuroscience at Washington University. More details at: https://www.humanconnectome.org/study/hcp-young-adult/document/hcp-citations.

## Footnotes

* Current versions of the toolboxes are publicly available on: https://users.fmrib.ox.ac.uk/~moisesf/cudimot/index.html https://users.fmrib.ox.ac.uk/~moisesf/Probtrackx_GPU/index.html

† A single CPU thread can use asynchronous CUDA commands (non-blocking) and send several tasks to different GPU streams; however, in our framework each streamline set needs a blocking post-processing step and therefore we need to use different CPU threads. We use OpenMP for creating multiple CPU threads.

‡ Note the almost sixfold difference between fitting NODDI-Bingham vs. NODDI-Watson using the Matlab toolbox, despite the higher complexity of NODDI-Bingham. This is due to the inefficient approximation of the confluent hypergeometric function of a scalar argument used in the NODDI-Watson Matlab implementation. For this reason, the comparisons in performance gains with cuDIMOT are more meaningful in the NODDI-Bingham case.

